# The MyMOMA domain of MYO19 encodes for distinct Miro-dependent and Miro-independent mechanisms of interaction with mitochondrial membranes

**DOI:** 10.1101/602938

**Authors:** Jennifer L. Bocanegra, Barbara M. Fujita, Natalie R. Melton, J. Matthew Cowan, Elizabeth L. Schinski, Tigist Y. Tamir, M. Ben Major, Omar A. Quintero

## Abstract

MYO19 interacts with mitochondria through a C-terminal membrane association domain (MyMOMA). The specific mechanisms for localization of MYO19 to mitochondria are poorly understood. Using new promiscuous biotinylation data in combination with existing affinity-capture databases, we have identified a number of putative MYO19-interacting proteins. We chose to further explore the interaction between MYO19 and the mitochondrial GTPase Miro2 by expressing mchr-Miro2 in combination with GFP-tagged fragments of the MyMOMA domain and assaying for recruitment of MYO19-GFP to mitochondria. Co-expression of MYO19^898-970^-GFP with mchr-Miro2 enhanced MYO19^898-970^-GFP localization to mitochondria. Mislocalizing Miro2 to filopodial tips or the cytosolic face of the nuclear envelope did not recruit MYO19^898-970^-GFP to either location. To address the kinetics of the Miro2/MYO19 interaction, we used FRAP analysis and permeabilization-activated reduction in fluorescence (PARF) analysis. MyMOMA constructs containing a putative membrane insertion motif but lacking the Miro2-interacting region displayed slow exchange kinetics. MYO19^898-970^-GFP, which does not include the membrane-insertion motif, displayed rapid exchange kinetics, suggesting that the MYO19 interacting with Miro2 has higher mobility than MYO19 inserted into the mitochondrial outer membrane. Mutation of well-conserved, charged residues within MYO19 or within the switch I and II regions of Miro2 abolished the enhancement of MYO19^898-970^-GFP localization in cells ectopically expressing mchr-Miro2. Additionally, expressing mutant versions of Miro2 thought to represent particular nucleotide states indicated that the enhancement of MYO19^898-970^-GFP localization is dependent on Miro2 nucleotide state. Taken together, these data suggest that membrane-inserted MYO19 is part of a larger complex, and that Miro2 plays a role in integration of actin- and microtubule-based mitochondrial activities.

## Introduction

The mitochondrial network serves multiple roles central to cellular physiology in addition to ATP homeostasis. Mitochondria are also important reservoirs for buffering intracellular calcium, regulation of reactive oxygen species, lipid synthesis, and apoptosis. [Friedman and Nunnari 2014]. Depending on the cell type and cell state, the geometry of the mitochondrial network can vary from a reticular arrangement of long threads, to a large number of smaller vesicular structures [Collins et al. 2002; Collins and Bootman 2003; Griparic and van der Bliek 2001]. The arrangement of the network is regulated by a balance of fission and fusion events [Mitra 2013], and the location of mitochondrial components within cells involves active positioning of these organelles in response to local need.

Mitochondrial interactions with the cytoskeleton play key roles in many of the processes related to mitochondrial dynamics and positioning. Loss of specific intermediate filament functions through either mRNA-knockdown, gene-knock-out, or mutation results in changes to mitochondrial motility, changes in mitochondrial distribution, or changes in metabolic activity in a number of different cell types [Chernoivanenko et al. 2015; Nekrasova et al. 2011; Schwarz and Leube 2016]. Much of our understanding of motor-protein-based transport comes from the study of mitochondrial motility in neurons, where long-range transport is thought to be mediated by interactions of mitochondria with microtubule based motors [Saxton and Hollenbeck 2012; Schwarz 2013]. There is a growing body of evidence indicating that microtubules and their associated proteins also influence fission and fusion dynamics [Mehta et al. 2019] [Perdiz et al. 2017]. Actin filaments and their associated proteins also contribute to the fission/fusion balance in cells [DuBoff et al. 2012; Hatch et al. 2016; Ji et al. 2015; Korobova et al. 2014; Korobova et al. 2013; Manor et al. 2015], as well as providing an additional system for motor-based transport through actin-myosin interactions [Kruppa et al. 2018; Pathak et al. 2010; Rohn et al. 2014]. It has recently been shown that Arp2/3-mediated actin filament clouds transiently assemble onto a large fraction (∼1/4) of the cellular mitochondria, promoting fission followed by fusion events once the cloud dissipates [Moore et al. 2016]. These clouds have been observed to cycle throughout the cytoplasm, interacting with the entirety of the mitochondrial network in approximately 15 minutes, thereby promoting the mixing of mitochondrial contents and prevention of large, inappropriately hyperfused mitochondrial arrangements.

A growing body of evidence suggests that the small GTPases Miro1 and Miro2 may be central signaling hubs that influence multiple cytoskeletal systems, having the potential to coordinate microtubule-based mitochondrial motility [Glater et al. 2006; MacAskill et al. 2009a] and fission/fusion events [Misko et al. 2010]. Miro1 and Miro2 are inserted into the mitochondrial outer membrane (MOM) through a C-terminal α-helix [Fransson et al. 2003]. The N-terminal 600 amino acids (approximately) face the cytoplasm, and consist of an N-terminal GTPase domain, two calcium binding EF hand motifs, and a second C-terminal GTPase domain [Fransson et al. 2003; Guo et al. 2005; Klosowiak et al. 2013] (domain structure illustrated in Supplemental Figure 1). Some reports indicate that Miro activities are dependent on calcium concentration [Macaskill et al. 2009b; Saotome et al. 2008; Wang and Schwarz 2009] and/or on the nucleotide state of the Miro GTPase domains [MacAskill et al. 2009a]. Although Miro1 and Miro2 proteins contain approximately 60% sequence homology [Fransson et al. 2003], differences exist in the cellular functions of the two paralogs and in their abilities to interact with other protein partners [Fransson et al. 2006; Lopez-Domenech et al. 2018; Lopez-Domenech et al. 2016; MacAskill et al. 2009a].

Recent reports published while we were completing these studies have also implicated Miro in actin-based mitochondrial functions. Mouse embryonic fibroblasts (MEF) Miro2 knockouts show decreased levels of the unconventional myosin, MYO19. MEFs with Miro1 and Miro2 knocked out show even lower levels of MYO19 [Lopez-Domenech et al. 2018]. HeLa cells with Miro1 and Miro2 knockdown showed a similar, low MYO19 phenotype [Oeding et al. 2018]. In both studies, decreases in cellular MYO19 levels was attributed to proteasome activity degrading MYO19 that was not mitochondria-associated [Lopez-Domenech et al. 2018; Oeding et al. 2018]. Additionally, the most C-terminal 72 amino acids of the MYO19 tail was shown to bind to the N-terminal GTPase domain of Miro1 *in vitro* [Oeding et al. 2018], potentially providing a mechanism for mitochondrial association and proteasome protection. Interestingly, ectopic expression of MYO19 in Miro double-knockout MEFs still resulted in mitochondria-associated MYO19, even in the absence of Miro proteins [Lopez-Domenech et al. 2018]. These data suggest that MYO19 contains a second mechanism of mitochondria association and that mitochondria-associated MYO19 may be less susceptible to proteasome activity.

MYO19 is a mitochondria-associated unconventional myosin [Quintero et al. 2009]. The N-terminal motor domain contains a high duty ratio [Lu et al. 2014], actin-activated ATPase capable of generating plus-end directed movements on actin filaments, *in vitro* [Adikes et al. 2013]. The mitochondria-binding, C-terminal MyMOMA domain contains a putative, monotopically inserted α-helix [Hawthorne et al. 2016; Shneyer et al. 2016] (domain structure illustrated in Supplemental Figure 1). The membrane-specificity of binding can be shifted to the endoplasmic reticulum by mutating two well-conserved basic residues located in close proximity to the monotopic insertion [Hawthorne et al. 2016; Shneyer et al. 2016]. However, ER-associated, GFP-tagged MYO19 constructs display faster FRAP and mobility kinetics than mitochondria-associated constructs [Hawthorne et al. 2016], suggesting that components of the mitochondrial outer membrane not present in ER membranes influence the mobility of MYO19. In some cell types, low-level ectopic expression of full-length MYO19 leads to a gain-of-function phenotype where the vast majority of the cellular mitochondrial network is motile [Quintero et al. 2009]. In other cell types, both endogenous MYO19 and ectopically expressed, fluorescently-tagged MYO19 have also been shown to localize with mitochondria at the tips of starvation-induced filopodia [Shneyer et al. 2017]. Loss of MYO19 function leads to a decreased ability to generate starvation-induced filopodia [Shneyer et al. 2016]. MYO19 knockdown also leads to two distinct defects tied to cell division [Rohn et al. 2014]: an increased failure rate at cytokinesis, and (for cells that proceed through cytokinesis) an unequal distribution of mitochondria in the resulting daughter cells.

Based on previously published data, we hypothesized that the MyMOMA domain of MYO19 encoded for multiple distinct mechanisms of interaction with the mitochondrial outer membrane, and that at least one mechanism was not strongly influenced by the presence of Miro proteins. We chose to identify potential MYO19 interactors using proteomics approaches and to investigate MYO19/Miro2 interactions using cell based, quantitative imaging approaches. Here we demonstrate that the Miro2/MYO19 interaction is potentially dependent on the nucleotide state of the N-terminal GTPase domain and that the kinetics of the Miro2/MYO19 interaction are faster-exchanging than those of the MYO19 membrane-inserted α-helix/MOM interaction. We propose a mechanism by which MYO19/Miro2 interactions would facilitate membrane-insertion of the monotopic α-helix, providing a means of regulating the levels of MYO19 on the mitochondrial outer membrane through Miro-mediated signaling pathways.

## Results and Discussion

### Interaction-analyses reveal potential MYO19 binding partners

MYO19 interacts with the mitochondrial outer membrane through the MyMOMA domain [Quintero et al. 2009], a sequence of ∼150 amino acids containing many basic residues [Hawthorne et al. 2016] and a putative monotopic membrane insertion [Shneyer et al. 2016]. The MyMOMA domain lacks sequence homology with other known proteins, making prediction of interacting partners challenging. In order to identify potential MYO19 binding partners, we generated a plasmid containing the promiscuous biotin ligase, BioID2 [Kim et al. 2016], attached to the C-terminus of the MyMOMA domain (amino acids 824-970) of human MYO19 (Supplemental Figure 1). We included a 30 amino acid linker sequence between BioID2 and MyMOMA to minimize any steric hindrance that might impact the ability of BioID2 to biotinylate neighboring proteins [Kim et al. 2016]. MYO19^824-970^-BioID2 localizes to mitochondria in HeLa cells stably expressing the construct, and when incubated with biotin, ALEXA350 streptavidin staining of mitochondria increased by approximately 40%, compared to the parent HeLa cells not expressing MYO19^824-970^-BioID2 (p<0.005, Figure 1A and 1B). Western blots from lysates of MYO19^824-970^-BioID2 expressing cells show more bands than control cells (Figure 1C), also indicating enhanced biotinylation, although to a lower level than in comparison with other cell lines and constructs under similar conditions [Kim et al. 2016; Roux et al. 2012]. We chose wild type HeLa cells as a control condition as we were assaying for enhanced biotinylation of proteins by BioID2 targeted to the mitochondrial membrane via the MyMOMA domain. As the enhancement in biotinylation was reproducible but not robust, using samples generated by expressing cytosolic BioID2 as a negative control had the potential of eliminating some bona fide candidate proteins from our data set: such as low-abundance proteins or proteins where cytosolic cellular pools exchange with mitochondrial pools. Although this approach might lead to some potential false positives, combining our proximity-labeling dataset with datasets using other experimental approaches to identify protein-protein interactions, we could identify the most likely candidate interacting proteins.

**Figure 1.**
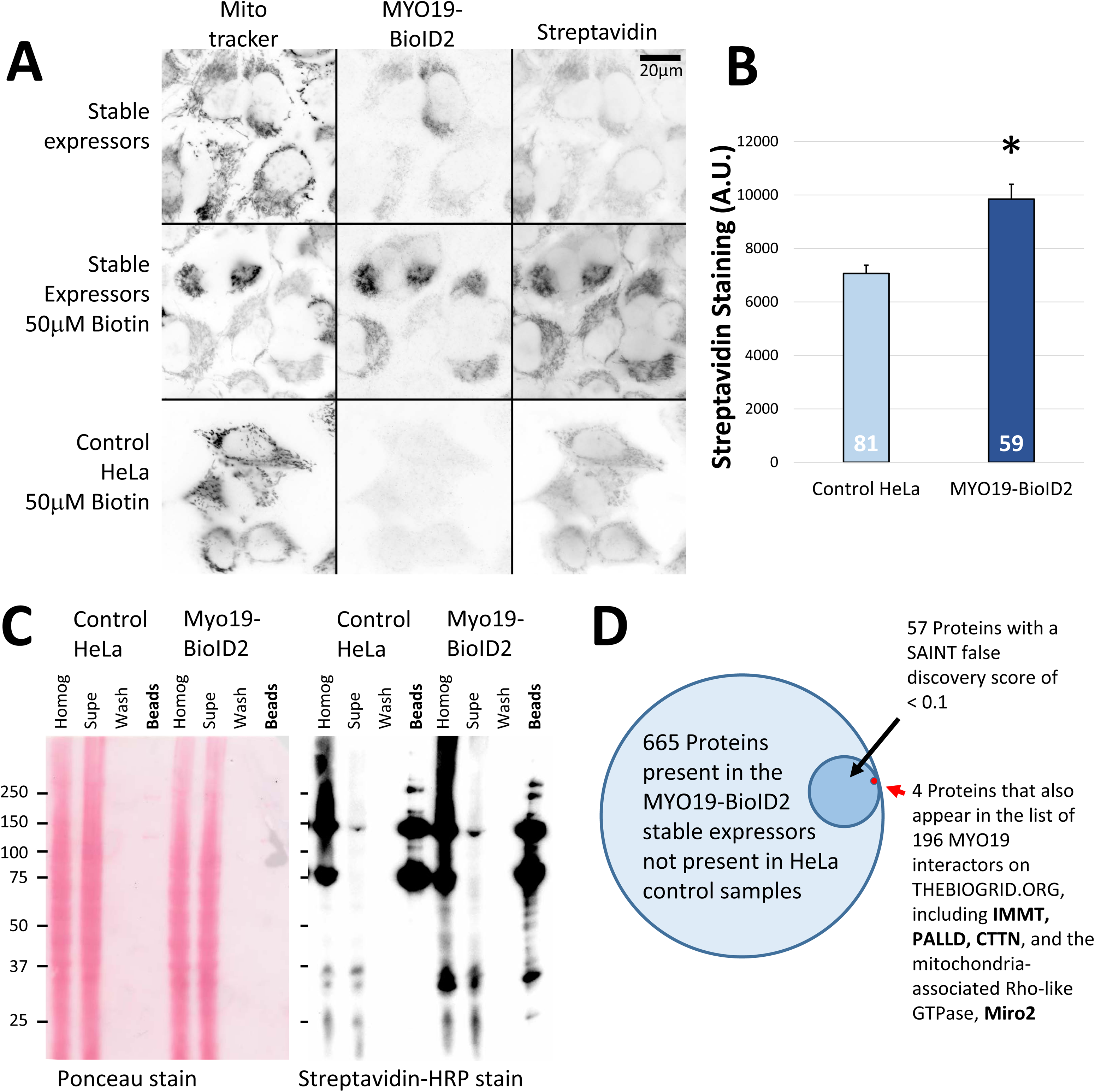
Putative MYO19 interactors can be identified using proteomic approaches. **A)** Cells stably expressing myc-MYO19^824-970^-BioID2 were generated via lentiviral infection, and showed biotinylation of mitochondrial proteins following overnight incubation with 50μM biotin. Expression of MYO19^824-970^-BioID2 was visualized via α-myc immunostaining, and biotinylation via ALEXA350-streptavidin staining. Each fluorescence channel was acquired under the same settings and displayed using the same scaling. The scale bar represents 20 μm. **B)** Quantification of ALEXA350-streptavidin staining indicates that MYO19^824-970^-BioID2 HeLa cells showed a ∼40% increase in biotinylation of mitochondrial-enriched regions compared to control HeLa cells (p<0.005, t-test). The number at the base of the bar is the number of cells (n) analyzed. **C)** HRP-streptavidin blot demonstrating increased biotinylation in HeLa cells stably expressing MYO19^824-970^-BioID2. Ponceau staining (left) demonstrates that the samples were loaded for equal cell number, and streptavidin-HRP staining (right) of the affinity bead sample lane from the BioID2 expressing cells shows more bands and darker labeling. **D)** Proteins from three sets of biotin-exposed cells stably expressing MYO19^824-970^-BioID2 were identified via mass spectroscopy. Of the 1420 proteins identified across all samples (including biotin-exposed, control HeLa cells), 665 proteins were not observed in control samples. 57 of those proteins had a SAINT false discovery rate < 0.1. Of those proteins, four were cross-listed in BIOGRID data for MYO19 interactors previously identified by affinity-capture mass spectrometry.

Upon establishing proper localization of the MYO19^824-970^-BioID2 construct and increased biotinylation in HeLa cells, we used mass spectrometry-based proteomics to identify the suite of biotinylated proteins from BioID2-MYO19^824-970^ expressing cells, compared to parental HeLa cells in biological triplicate. Multiple approaches can be taken to analyze the results of potential interactors identified by promiscuous biotinylation. MaxQuant analysis [Cox and Mann 2008] identified 3 low-confidence hits and 32 high-confidence hits based on label-free-intensity quantitation as well as peptide count (Supplemental Table 2). We also used the Significance Analysis of Interactome (SAINT) method to identify potential MYO19 interactors [Choi et al. 2011]. This scoring system not only accounts for whether a protein is present in the BioID2 sample, but whether it is significantly more enriched in the experimental versus the control sample. SAINT analysis of whole-cell lysates identified 57 biotinylated proteins that had a SAINT basal false discovery rate (BFDR) score less than 10% (Supplemental Table 2).

To determine which proteins within the list of MYO19-proximal proteins may interact with MYO19, we cross-referenced our biotinylation data with published protein-protein interaction data available at THEBIOGRID.org, an online repository of peer-reviewed, published, protein-protein interaction data [Stark et al. 2006]. In the case of MYO19, three separate studies using affinity-capture in combination with mass spectrometry have identified a total of 198 MYO19 interactors [Boldt et al. 2016; Choudhury et al. 2017; Giurato et al. 2018; Hein et al. 2015; Huttlin et al. 2017; Huttlin et al. 2015; Khanna et al. 2018] (Supplemental Table 2). Cross-referencing the MaxQuant list of high-confidence (32) and low-confidence (3) hits with data from THEBIOGRID.org resulted in an overlap of 6 targets: palladin (PALLD), mitofilin (IMMT), filamin A (FLNA), filamin B (FLNB), voltage-dependent anion-selective channel protein 2 (VDAC2), and neuroblast differentiation-associated protein (AHNAK). Of the 57 proteins with a SAINT BFDR < 0.1, five proteins overlapped with THEBIOGRID.org data (Figure 1D): MYO19 itself (BFDR < 0.0001), mitofilin (BFDR < 0.0001), palladin (BFDR < 0.0001), cortactin (CTTN, BFDR < 0.0001), and Miro2 (RHOT2, BFDR < 0.0892). Comparing the two sets, cortactin and Miro2 were identified in the MaxQuant data, but not identified as high- or low-confidence hits. Filamin A, filamin B, VDAC2, and AHNAK were not present in the SAINT list of 665 proteins not observed in the control samples. Oeding and colleagues identified 41 proteins biotinylated by BioID-MYO19^824-970^ [Oeding et al. 2018]. Of those proteins, 10 overlap with our SAINT list, four with a BFDR < 0.1, including Miro2 (Supplemental Table 2). Although both sets of data result from stable expression in HeLa cells of a MyMOMA domain construct, the Oeding construct was N-terminally tagged with a different promiscuous biotin ligase [Roux et al. 2012] and a short, two amino-acid linker. Our construct contained a smaller, more enzymatically active biotin ligase [Kim et al. 2016] with a longer linker attached to the C-terminus of MYO19^824-970^. As different termini of the MyMOMA domain might be positioned differently in their cellular environment, tagging different ends of the MyMOMA domain could result in the identification of distinct sets of neighboring proteins, and linker length has been shown to influence biotinylation activity [Kim et al. 2016].

Based on these analyses, multiple proteins with functional links to the cytoskeleton and mitochondrial function have been identified. Mitofilin localizes to the inner membrane space [Odgren et al. 1996] and is associated with maintenance of cristae morphology [John et al. 2005; Zerbes et al. 2012]. The *Drosophila* homolog, MIC60, has been shown to regulate mitochondrial dynamics. Mutant flies lacking MIC60 show decreased Miro protein levels and altered mitochondrial dynamics [Tsai et al. 2017]. VDAC2 are mitochondrial outer membrane porins involved in ATP homeostasis [Colombini 2012; Saks et al. 1995], and in some instances may be regulated by interaction with microtubules [Ramos et al. 2019]. AHNAK is a large scaffolding protein with the ability to bind actin and other cellular components in a variety of cellular contexts [Davis et al. 2014], including muscle contraction [Haase et al. 2004], cancer [Shankar et al. 2010], and Schwann cell function [von Boxberg et al. 2014]. Filamins are actin crosslinking proteins involved in actin-based cellular activities via interactions with a large number of other protein partners [Nakamura et al. 2011]. It was recently reported that DRP1, a component of the mitochondrial fission machinery, and filamin A interact in response to hypoxic stress [Nishimura et al. 2018]. Palladin is an actin crosslinking protein originally characterized in the stress fibers of fibroblasts and cardiac myocytes [Parast and Otey 2000]. Palladin is a cytosolic Ig domain containing protein with multiple isoforms [Rachlin and Otey 2006], and has multiple roles in organizing the actin cytoskeleton in many structures and cell types [Goicoechea et al. 2008; McLane and Ligon 2015; Sun et al. 2017]. To date little evidence exists for interactions with mitochondria. Cortactin often localizes to the regions of dynamic actin, such as membrane ruffles and lamellipodia, where it mediates actin filament branching through interactions with the Arp2/3 complex [Schnoor et al. 2018]. A recent report has also shown a role for cortactin in the assembly of mitochondria-associated actin structures involved in mitochondrial network morphology and fission [Li et al. 2015].

Miro2, a member of the Ras superfamily of GTPases [Colicelli 2004; Jaffe and Hall 2005], is one of two paralogs that exist in the human genome (the other being Miro1) [Fransson et al. 2003]. Miro/dynein functional interactions have been identified in multiple cell types [Babic et al. 2015; Melkov et al. 2016; Morlino et al. 2014; Russo et al. 2009; van Spronsen et al. 2013], as have Miro/kinesin interactions [Glater et al. 2006] via Milton/Trak/GRIF proteins [Guo et al. 2005; MacAskill et al. 2009a; Stowers et al. 2002]. Miro proteins have also been shown to be a component of protein clusters implicated in cristae maintenance and ER-mitochondria contact site maintenance [Modi et al. 2019].

### Ectopic expression of Miro2 enhances localization of fast-exchanging MYO19^898-970^ to mitochondria

Recently, work from two other research groups have implicated Miro proteins in the stability of MYO19 protein levels in the cell [Lopez-Domenech et al. 2018] and in mitochondria interactions [Oeding et al. 2018]. Lopez-Domenech and colleagues were able to immunoprecipitate endogenous MYO19 together with Miro2 [Lopez-Domenech et al. 2018], and Oeding and colleagues were able to show that bacterially-expressed MYO19^898-970^ and Miro1^1-592^ interacted in an *in vitro* binding assay [Oeding et al. 2018]. Interestingly, although Miro proteins were hypothesized to serve as receptors for MYO19 on mitochondria, ectopic expression of MYO19 constructs lacking the putative Miro-interacting region are able to localize to mitochondria [Hawthorne et al. 2016; Shneyer et al. 2016], likely through a distinct membrane-inserted monotopic α-helix. Additionally, ectopic expression of MYO19 constructs in Miro doubled-knockout MEFs still results in mitochondrial localization of those constructs [Lopez-Domenech et al. 2018].

As the Miro-interacting region of MYO19 does not include the putative monotopic membrane insertion [Shneyer et al. 2016], we chose to assess whether the combination of the Miro-interacting region and the monotopic membrane insertion were required for Miro-enhanced mitochondrial localization. We compared the mitochondrial localization phenotype observed for MYO19^853-935^-GFP (partial deletion of the putative Miro-interacting domain) and MYO19^860-890^-GFP (complete deletion of the Miro-interacting domain) expressed in the presence of mchr-Miro2 and compared that to the phenotype of MYO19^898-970^-GFP expressed in the presence of mchr-Miro2 (see Supplemental Figure 1 for a construct map). We examined the enhancement of mitochondrial localization by measuring the ratio of mitochondrial GFP fluorescence to cytosolic GFP fluorescence [Hawthorne et al. 2016] (mito/cyto ratio, Figure 2A). We also examined the penetrance of the phenotype by quantifying the fraction of cells in a population showing diffuse cytoplasmic GFP staining, strong mitochondrial staining, or intermediate staining (Figure 2B, Supplemental Figure 2A).

**Figure 2.**
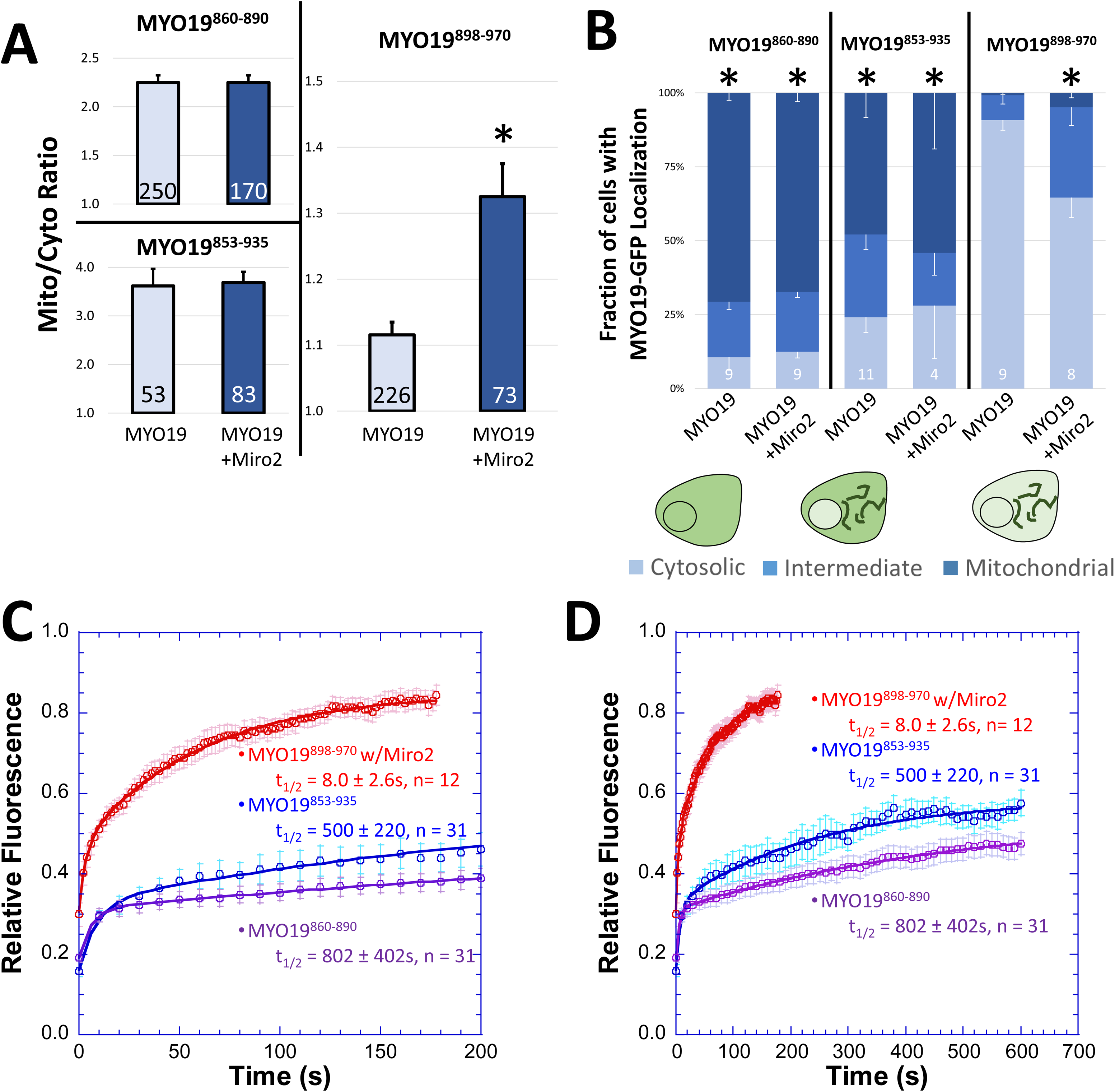
MYO19^898-970^ localizes to mitochondria in the presence of Miro2 and has more rapid exchange kinetics than constructs containing the putative membrane-insertion motif. **A)** Mito/cyto ratio was used to quantify the enhancement of mitochondrial localization of MYO19-GFP constructs in the presence of mchr-Miro2. While coexpression of mchr-Miro2 with MYO19^898-970^-GFP enhanced the GFP-localization on mitochondria, mitochondrial localization of MYO19^853-935^-GFP and MYO19^860-890^-GFP were not enhanced by the coexpression of mchr-Miro2 (*p < 0.001, t-test between MYO19^898-970^-GFP and MYO19^898-970^-GFP coexpressed with mchr-Miro2). **B)** Coexpression of mchr-Miro2 decreased the fraction of the population of MYO19^898-970^-GFP expressing cells displaying a cytosolic staining pattern for GFP, but expression of mchr-Miro2 did not alter the population of MYO19^853-935^-GFP or MYO19^860-890^-GFP expressing cells with a cytosolic GFP pattern, further implicating Miro2 in localizing MYO19 to mitochondria (* p <0.05, compared to MYO19^898-970^-GFP, Tukey analysis). **C)** FRAP recovery of MYO19^853-935^-GFP and MYO19^860-890^-GFP expressing cells is relatively slow with a large immobile fraction, when compared to MYO19^898-970^-GFP in cells expressing mchr-Miro2, indicating that the mechanism of interaction with mitochondria for these two regions of MYO19 differs. **D)** Same data as in (C), but plotted on a different time scale to show the complete time course for the longer recovery kinetics. FRAP kinetic parameters are listed in Table 1.

While ectopic expression of mchr-Miro2 enhanced MYO19^898-970^-GFP localization to mitochondria (as determined by the mito/cyto ratio) and increased the fraction of cells with an intermediate or strong mitochondrial localization pattern (as determined by population analysis), mchr-Miro2 coexpression did not enhance the mito/cyto ratio for MYO19^853-935^-GFP or MYO19^860-890^-GFP (Figure 2A), nor did it alter the population of cells displaying a mitochondria-localized phenotype (Figure 2B). Additionally, it is worth noting that the mito/cyto ratio for cells expressing MYO19^898-970^ in the presence of mchr-Miro2 (∼1.3) is lower than the mito/cyto ratio for cells expressing either of the other two constructs which contain the membrane insertion domain (greater than 2, Figure 2A). Also, the fraction of cells displaying some mitochondrial localization for cells expressing MYO19^898-970^ in the presence of mchr-Miro2 (∼35%) is lower than for cells expressing MYO19^853-935^-GFP or MYO19 ^860-890^-GFP (greater than 70%, Figure 2B). These data suggest that membrane-insertion domain may have a stronger influence on mitochondrial association than does Miro interaction, and that two different mechanisms exist within the MyMOMA domain for binding to mitochondria outer membrane components.

We hypothesized that the different mechanisms of mitochondrial outer membrane-binding encoded by different regions of the MyMOMA domain would have different affinities, and that those differences could be revealed by examining the kinetics of interaction between the mitochondrial outer membrane and each region of the MyMOMA domain. We compared protein mobility and the equilibrium kinetics of exchange through fluorescence recovery after photobleaching analysis (FRAP) [Lippincott-Schwartz et al. 2003; Reits and Neefjes 2001]. For these experiments, we chose to favor Miro2 interactions for the MYO19^898-970^-GFP construct by coexpressing mchr-Miro2. The relative fluorescence of the regions of interest remained steady prior to bleaching (Supplemental Figure 2B). FRAP recovery of MYO19^898-970^-GFP was approximately 30-fold faster than the recovery for MYO19^853-935^-GFP or MYO19^860-890^ (Figures 2C and 2D, Table 1). Additionally, the immobile fraction for MYO19^898-970^-GFP was approximately 15-fold smaller than that calculated for MYO19^853-935^-GFP or MYO19^860-890^-GFP (Figures 2C and 2D, Table 1). We also examined the non-equilibrium, relative dissociation kinetics of GFP-tagged MYO19 truncations via permeabilization activated reduction in fluorescence kinetics (PARF) [Singh et al. 2016]. PARF analysis indicated that the off-rate kinetics for MYO19^898-970^-GFP were faster, resulting in more complete loss of fluorescence over the time course of the experiment than what was observed for MYO19^853-935^-GFP (Supplemental Figure 2C, Table 2). While the PARF half-life for loss of MYO19^898-970^-GFP fluorescence was ∼80s and the calculated immobile fraction was∼33%, MYO19^853-935^-GFP did not drop below ∼70% of its initial fluorescence, so a half-life could not be calculated from conditions with such a large immobile fraction. Taken together, the association data and kinetic data indicate that two mechanisms for mitochondrial outer membrane interaction are encoded by different regions of the MyMOMA domain. The MYO19/Miro2 interaction mediated by amino acids 898-970 is a faster-exchanging, higher mobility interaction than the interaction mediated by the membrane-inserting α-helix motif, suggesting a lower affinity interaction between MYO19 and Miro compared to MYO19 that has been inserted into the mitochondrial outer membrane.

**Table 1:** FRAP kinetic analysis for proteins used in this study.

| Construct | $t_{1/2}$ (s) | Immobile | R | n |
| --- | --- | --- | --- | --- |
| MYO19<sup>853-935</sup>-GFP | 599±220 | 0.47±0.03 | 0.9935 | 31 |
| MYO19<sup>860-890</sup>-GFP | 802±402 | 0.45±0.02 | 0.99741 | 31 |
| MYO19<sup>898-970</sup>-GFP/mchr-Miro2 wt* | 9.2±1.7 | 0.03±0.02 | 0.9986 | 39 |
| MYO19<sup>898-970</sup>-GFP **mut.**/mchr-Miro2 wt | 1.5±0.3 | 0.04±0.06 | 0.9925 | 21 |
| MYO19<sup>898-970</sup>-GFP/mchr-Miro2 **mut.** | 9.0±1.6 | 0.06±0.01 | 0.9981 | 19 |
| MYO19<sup>898-970</sup>-GFP/mchr-Miro2 **A13V** | 8.0±0.9 | 0.14±0.02 | 0.9995 | 18 |
| MYO19<sup>898-970</sup>-GFP/mchr-Miro2 **T18N** | 17±4.0 | 0.16±0.02 | 0.9994 | 20 |

\*combination of all MYO19<sup>898-970</sup>-GFP/mchr-Miro2 samples across all trials

**Table 2:** PARF kinetic analysis for proteins used in this study.

| Construct | $t_{1/2}$ (s) | Immobile | R | n |
| --- | --- | --- | --- | --- |
| MYO19<sup>853-935</sup>-GFP | - | 0.69±0.04 | 0.9531 | 49 |
| MYO19<sup>898-970</sup>-GFP | 13±4.3 | 0.12±0.03 | 0.9926 | 16 |
| MYO19<sup>898-970</sup>-GFP/mchr-Miro2 wt | 83±12 | 0.33±0.02 | 0.9976 | 44 |

Oeding et al. previously reported that mislocalization of Miro1 to endosomes resulted in correlation between the MYO19^898-970^ signal and endosomal markers, which they interpreted as recruitment of MYO19^898-970^ to endosomes [Oeding et al. 2018]. To address whether similar recruitment occurred with Miro2, we relocalized mchr-Miro2 lacking the mitochondrial outer membrane insertion sequence (mchr-Miro2ΔTM) to filopodial tips by coexpressing it with a chimeric construct consisting of an α-mcherry nanobody [Fridy et al. 2014] attached to the C-terminus of a MYO10 fragment capable of localizing to filopodial tips [Berg and Cheney 2002], MYO10-HMM-nanotrap^red^, which was based on a previously-published construct containing an α-GFP nanobody [Bird et al. 2017]. Coexpression of MYO10-HMM-nanotrap^red^, mchr-Miro2ΔTM, and MYO19^898-970^-GFP resulted in the concentration of mcherry signal at the filopodial tips, but not GFP signal, as indicated by the ratio of filopodial tip intensity to cytoplasmic intensity (Supplemental Figure 3A). Coexpression of MYO10-HMM-nanotrap^green^ (contains an α-GFP nanobody instead of an α-mcherry nanobody), mchr-Miro2ΔTM, and MYO19^898-970^-GFP led to concentration of GFP signal at the filopodial tip, but not mcherry signal (Supplemental Figure 3A). To validate whether cooccurrence and localization of two labeled constructs to filopodial tips was possible in this system, we coexpressed MYO10-HMM-nanotrap^green^, mchr-Miro2ΔTM, and a GFP-tagged nanobody against mcherry [Fridy et al. 2014], GFP-nanotrap^red^. In this instance, both GFP and mcherry signal concentrated at filopodial tips, as indicated by a filopodial tip intensity/cell body intensity ratio greater than 1 in both channels (Supplemental Figure 3A). To ensure that the geometry of filopodia was not interfering with the ability of the constructs to interact, we also localized mchr-Miro2 to the cytosolic face of the nuclear envelope by replacing the mitochondrial localization sequence with the KASH domain from nesprin-2β [Luxton et al. 2010]. High levels of overexpression of the mchr-Miro2-KASH construct led to aggregates and improper localization of the construct, but low expressors showed nuclear envelope localization of the mcherry construct (Supplemental Figure 3B). Coexpression of mchr-Miro2-KASH with MYO19^898-970^-GFP did not show cooccurrence of mchr and GFP signal on the nuclear membrane, indicating that mchr-Miro2-KASH was not recruiting MYO19^898-970^ (Supplemental Figure 3B). As a control, coexpression of a GFP-nanobody [Rothbauer et al. 2008] targeted to the nuclear envelope via a KASH domain (mchr-nanotrap^green^-KASH) coexpressed with MYO19^898-970^-GFP did result in cooccurrence of the GFP signal and mchr signal on the nuclear membrane, indicating recruitment of the MYO19 construct to the nuclear envelope visualized as a ring of high-intensity fluorescence surrounding the nucleus in both channels (Supplemental Figure 3B). Oeding et. al demonstrated that Miro1 mislocalized to endosomes was capable of recruiting C-terminally tagged GFP-MYO19^898-970^, using a different mislocalization and analysis approach than the two approaches that we employed.

These data may indicate that functional differences exist in the ability of Miro1 and Miro2 to interact with binding partners. Additionally, differences in DNA dose during transfection could accentuate low-affinity interactions in cells expressing high protein levels. As previous researchers reported that overexpressing Miro constructs impacted mitochondrial morphology and cell function [Fransson et al. 2003; Liu et al. 2012; Russo et al. 2009], we carried out all of our expression experiments by transfecting low amounts of DNA and completing our observations within 18-30 hours after transfection. These conditions resulted in a transfection efficiency of approximately ∼15%. By performing immunocytochemistry on the subpopulation of transfected cells using an α-MYO19 antibody that did not recognize ectopically-expressed MYO19 constructs, we were able to verify that ectopic expression of either MYO19^898-970^-GFP and/or mchr-Miro2 reduced the endogenous expression of MYO19 by approximately 10-15%, compared to untransfected neighboring cells prepared under the same conditions (data not shown).

### Miro2-dependent localization of MYO19^898-970^ may be influenced by Miro2 nucleotide state

One mechanism by which small GTPases interact with their effector proteins is via the formation of a binding surface. When in the GTP-bound state, the small GTPase surface loops known as switch I and switch II are positioned in close proximity [Dvorsky and Ahmadian 2004; Vetter and Wittinghofer 2001]. We hypothesized that such a conformational change might mediate MYO19/Miro2 interactions. To address this possibility, we first examined the conservation of Miro2 sequences across seven vertebrate species (mouse, dingo, human, cow, zebrafish, chicken, and *Xenopus tropicalis*), looking for conservation within switch I and switch II (Supplemental Figure 4A). We identified a 6 amino-acid stretch in switch I, XXFPXX, where X represents well-conserved acidic residues. A similar stretch, SXAXQTXXX, was well conserved in switch II (Figure 3A). We had previously identified well conserved, basic residues in the MYO19 tail [Hawthorne et al. 2016], and hypothesized that interactions between a basic patch in MYO19^898-970^ (Supplemental Figure 4B) and an acidic patch on Miro2 would mediate this interaction (Figure 3A). To test this hypothesis, we generated MYO19^898-970^-GFP constructs where K^923^, R^927^, and K^928^ were mutated to alanine. When coexpressed with wild type mchr-Miro2, the mutant MYO19^898-970^-GFP did not show enhanced mitochondrial localization as determined by the GFP mito/cyto ratio (Figure 3B), nor did it increase the population of cells displaying an enhanced mitochondrial localization phenotype (Figure 3C).

**Figure 3.**
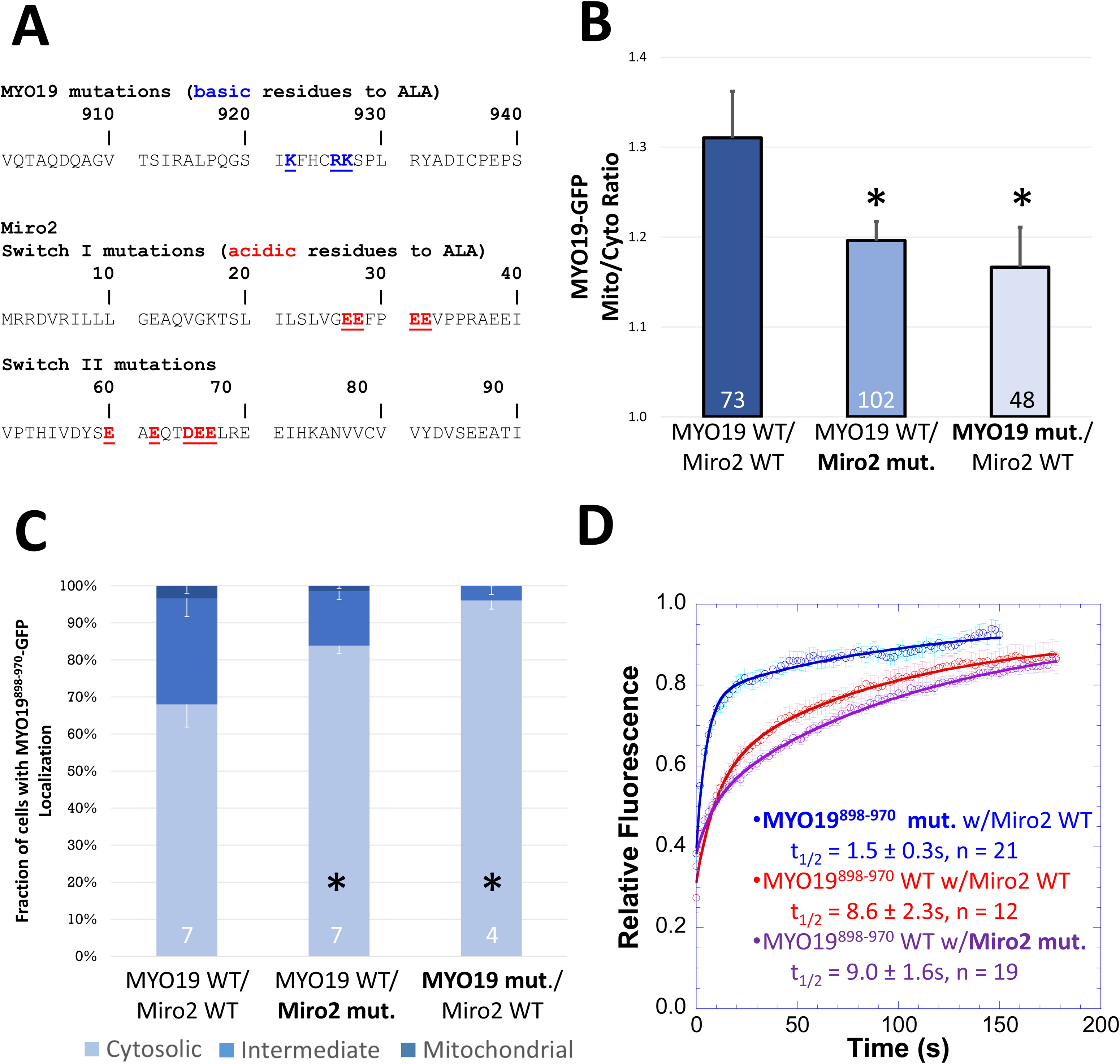
A subset of well-conserved, charged residues are essential for Miro2-dependent localization of MYO19^898-970^. **A)** Clustal Omega multiple sequence alignments of the MYO19 MyMOMA domain and the Miro2 N-terminal GTPase domain identified well-conserved, charged residues in each domain, which were mutated to alanine to determine their contribution to the MYO19/Miro2 interaction. **B)** Mutating either the basic residues in MYO19^898-970^-GFP to alanine or the acidic residues in mchr-Miro2 to alanine decreased the localization of MYO19^898-970^-GFP to mitochondria, as determined by the mito/cyto ratio (*p <0.05, Dunnett’s test versus MYO19^898-970^-GFP wild type coexpressed with mchr-Miro2 wild type). **C)** Mutations similarly decreased the fraction of cells displaying a cytosolic staining pattern for GFP, whether those mutations were to MYO19 or Miro2 (*p <0.05, Dunnett’s test versus MYO19^898-970^-GFP wild type coexpressed with mchr-Miro2 wild type). **D)** FRAP analysis of MYO19^898-970^-GFP mutant shows faster exchange kinetics than wild type, consistent with basic residues in the MYO19 mediating interactions with Miro2. The exchange kinetics for wild type MYO19^898-970^-GFP with the mutant mchr-Miro2 construct were similar to exchange kinetics with the wild type mchr-Miro2 wild type construct. This is likely due to endogenous Miro recruiting MYO19^898-970^-GFP. Multiple sequence alignments can be found in the Supplemental Figure 4. Numbers at the base of the bars indicate the number of replicates. Error bars represent standard error of the mean. FRAP kinetic parameters are listed in Table 1.

We also generated a mchr-Miro2 construct where 4 acidic residues in switch I and 5 acidic residues in switch II were mutated to alanine. When this construct was coexpressed with wild type MYO19^898-970^-GFP, the GFP-construct did not show enhanced GFP mito/cyto ratio, nor did the fraction of cells displaying a mitochondrial localization phenotype increase (Figure 3B and 3C). FRAP analysis of cells showing mitochondrial localization revealed a 3-fold faster half-life for cells expressing the mutant MYO19^898-970^-GFP and wild type mchr-Miro2, when compared to cells expressing wild type MYO19^898-970^-GFP and wild type mchr-Miro2 (Figure 3D, Table 1). The half-life of wild type MYO19^898-970^-GFP in cells expressing the mutant mchr-Miro2 construct (9.0±1.6s) was similar to that of cells expressing wild type MYO19^898-970^-GFP and wild type mchr-Miro2 (9.2±1.7s). This result is likely due to the fact that cells used in FRAP experiments were chosen for analysis only if they displayed strong mitochondrial localization, which was likely due to the presence of endogenously-expressed Miro1 and Miro2. These data suggest that a charge interaction mediates the interaction of MYO19^898-970^ with Miro2.

If the Miro2/MYO19 interaction is indeed mediated by the formation of an acidic patch on Miro2 in a manner that is dependent on the nucleotide state of the N-terminal GTPase [Fransson et al. 2006], then expressing mutant forms of Miro2 predicted to favor either the GTP-bound [Diekmann et al. 1991] or GDP-bound state [Farnsworth and Feig 1991] would influence the ability of the mchr-Miro2 to interact with MYO19^898-970^-GFP (Figure 4A). While expressing mchr-Miro2 A13V (GTP-bound state mimic) in combination with MYO19^898-970^-GFP enhanced the GFP mito/cyto ratio and the fraction of cells showing a mitochondrial localization phenotype at levels similar to wild type mchr-Miro2, expression of mchr-Miro2 T18N (GDP-bound state mimic) did not enhance either of those two properties (Figure 4B and 4C). FRAP analysis revealed similar equilibrium kinetics for MYO19^898-970^ expressed with mchr-Miro2 wild type, mchr-Miro2 A13V, or mchr-Miro2 T18N (Figure 4D, Table 1). As cells with strong mitochondrial localization of GFP construct were chosen for FRAP analysis, it is likely that the interaction being measured under these experimental conditions contains a contribution from endogenous Miro1 and Miro2, whose activity is not masked by the ectopic expression of mutant Miro constructs. This may have contributed to the similarity in FRAP kinetics between Myo19^898-970^ expressed in a wild type mchr-Miro2 background compared to the kinetics of the same protein expressed in the mchr-Miro2^T18N^, GDP-mimic background. Taken together, these data are consistent with an electrostatic interaction between MYO19 and Miro2 that can be regulated by the nucleotide-state of Miro2. Additionally, it also suggests that the nucleotide state of exogenously expressed wild type mchr-Miro2 is likely the GTP-bound state as the binding properties (as measured by mito/cyto ratio and by the penetrance of the mitochondrial-localization phenotype) and kinetic properties of the mchr-Miro2 A13V construct were very similar to those of the mchr-Miro2 wild type construct.

**Figure 4.**
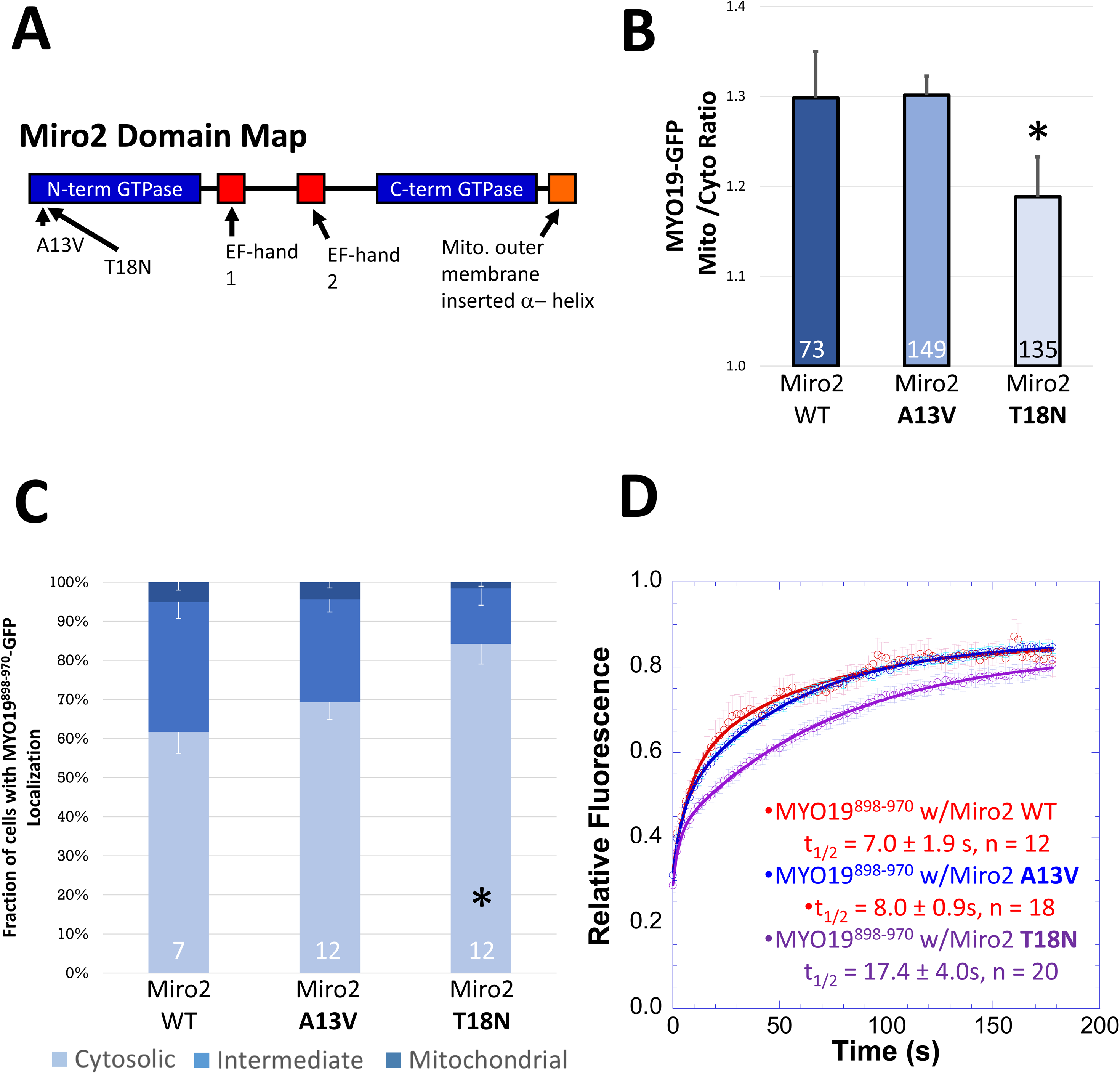
Miro2-dependent localization of MYO19^898-970^ may depend on the nucleotide state of Miro2. **A)** Point mutations made to the N-terminal domain of Miro2 are thought to result in the GTPase being in the GTP-bound (A13V) or GDP-bound (T18N) state. **B)** Expressing the putative GDP-bound mchr-Miro2 T18N mutant decreased the localization of MYO19^898-970^-GFP to mitochondria while the putative GTP-bound A13V mutant did not, as determined by the mito/cyto ratio (*p <0.05, Dunnett’s test versus MYO19^898-970^-GFP coexpressed with mchr-Miro2 wild type). **C)** Expression of mchr-Miro2 T18N increased the fraction of cells displaying a cytosolic staining pattern for GFP (*p <0.05, Dunnett’s test versus MYO19^898-970^-GFP coexpressed with mchr-Miro2 wild type). **D)** FRAP analysis of MYO19^898-970^-GFP with wild type or GTP-state mutants shows similar exchange kinetics of MYO19^898-970^-GFP for all three conditions. For the mchr-Miro2 T18N, this is likely due to endogenous Miro recruiting MYO19^898-970^-GFP. Numbers at the base of the bars indicate the number of replicates. Error bars represent standard error of the mean. FRAP kinetic parameters are listed in Table 1.

### Implications of a central signaling protein integrating microtubule-based and actin-based motor interactions with mitochondria

The available proteomics data suggests that MYO19 interacts with or is in close-proximity to a number of proteins involved in mitochondrial functions tied to cytoskeletal activities. Many of these interactions are with proteins that serve roles in assembling larger protein complexes (mitofilin, paladin, filamin, and AHNAK). Such interactions have the potential of contributing to the slower mobility and exchange dynamics of mitochondria-associated MYO19 [Reits and Neefjes 2001]. It is worth noting that when MYO19^853-935^-GFP was mislocalized to the ER, its FRAP kinetics were 10-fold faster than for the mitochondria-localized construct [Hawthorne et al. 2016], perhaps because additional, mitochondria-specific MYO19 interactors were not present in the ER. Similarly, the FRAP and mobility kinetics of the Miro-interacting domain of MYO19 were more rapid than those of the constructs containing the membrane-insertion motif, further supporting the hypothesis that different regions of the MyMOMA domain encode for the ability to interact with distinct binding partners. Further research will be required to more carefully examine the other components of mitochondria that interact with MYO19, and how those interactions influence MYO19 dynamics and mitochondrial biology.

We chose to focus on Miro2 initially as it has the potential for being a central regulator of multiple mitochondria/cytoskeleton interactions, including anchoring, fission, and transport. Taken together, our data and previous studies [Lopez-Domenech et al. 2018; Oeding et al. 2018] are elucidating a role for Miro1 and Miro2 in mediating the interaction between MYO19 and mitochondria. Lopez-Domenech and colleagues demonstrated that Miro-knockdown MEFs had significantly decreased levels of MYO19, and that in single-knockouts, Miro2 had a more drastic effect than Miro1 on MYO19 levels. Similarly, Oeding and colleagues demonstrated that cells treated with siRNA for Miro1 and Miro2 also displayed a significant decrease in MYO19 protein levels. Both groups also reported that Miro double knockdown or knockout resulted in more rapid loss of MYO19 protein than in wild type cells. According to Lopez-Domenech and colleagues, this was due to proteasome-mediated degradation [Lopez-Domenech et al. 2018]. Interestingly, fluorescently-tagged constructs lacking part of the Miro-interacting domain of MYO19 [Hawthorne et al. 2016] or all of the Miro interacting domain of MYO19 [Shneyer et al. 2016] will still localize to the mitochondrial outer membrane, so long as they contain an intact monotopic membrane inserting α-helix. Additionally, MYO19-GFP expressed in Miro1/2 double-knockout cells is able to colocalize with mitochondria [Lopez-Domenech et al. 2018].

The differences between these observations can be explained based on the kinetics of the proteasome degradation machinery, the concentration of available MYO19, and the kinetics of MYO19 membrane insertion. Such an explanation also provides a hypothesis for how Miro activity could modulate MYO19 function. If Miro proteins are providing a fast-exchanging binding site on the mitochondrial membrane for MYO19, then when MYO19 is in proximity of the mitochondrial membrane, rare interactions (like monotopic insertion) are favored over degradation pathways. In this way, initial Miro-dependent recruitment of MYO19 to the mitochondrial outer membrane prolongs the lifetime of MYO19 in cells by promoting additional MYO19/MOM interactions that do not require Miro. In cells expressing normal levels of MYO19 and Miro proteins, the facilitation of MYO19 membrane insertion by Miro is sufficient to overcome the proteasome-mediated degradation pathways (Figure 5A). If membrane-inserted MYO19 has a longer half-life than cytosolic MYO19, then the pool of mitochondrial outer membrane-associated MYO19 is maintained. In the case of cells lacking Miro, cytosolic MYO19 is degraded more quickly than it can be efficiently membrane-inserted, leading to decreased levels of mitochondria associated MYO19 as well as an overall decrease in cellular levels of MYO19 (Figure 5B). The same would be true if the nucleotide state of Miro regulated its ability to interact with MYO19 (Figure 5C). In the case of constructs lacking the Miro-interacting domain, ectopic expression leading to higher MYO19 concentrations than under endogenous conditions would lead to more occurrences of rare membrane insertion events. If MYO19 levels are higher than what proteasome-mediated degradation pathways can process, then an appreciable number of membrane-insertion events may still occur. It may also be the case that once membrane-inserted, MYO19/Miro interactions may protect MYO19 from degradation (Figure 5D). This interpretation is consistent with the observation that the turnover of exogenously-expressed, Halo-tagged MYO19 is faster in Miro1/Miro2 double knockdown cells than in wild type HeLa cells [Oeding et al. 2018]. Additionally, Miro1 levels decrease more rapidly than MYO19 levels in FCCP-treated cells [Lopez-Domenech et al. 2018], suggesting that Miro1 is protecting MYO19 from degradation, that Miro1 is more readily degraded than MYO19, or both. Interestingly, inadvertent protection mechanisms may have actually facilitated the discovery of the monotopic membrane-insertion domain by Scheyer, Hawthorne, and their colleagues. Oeding et al. reported that C-terminal tagged MYO19 constructs showed slower degradation that N-terminally tagged MYO19 constructs [Oeding et al. 2018]. Both Hawthorne et al. and Scheyer et al. used C-terminally tagged MYO19 constructs in their studies, and since these constructs were overexpressed and may have been slower to degrade, it would have been easier to identify fluorescence localization to mitochondria even without Miro-facilitated membrane association.

**Figure 5.**
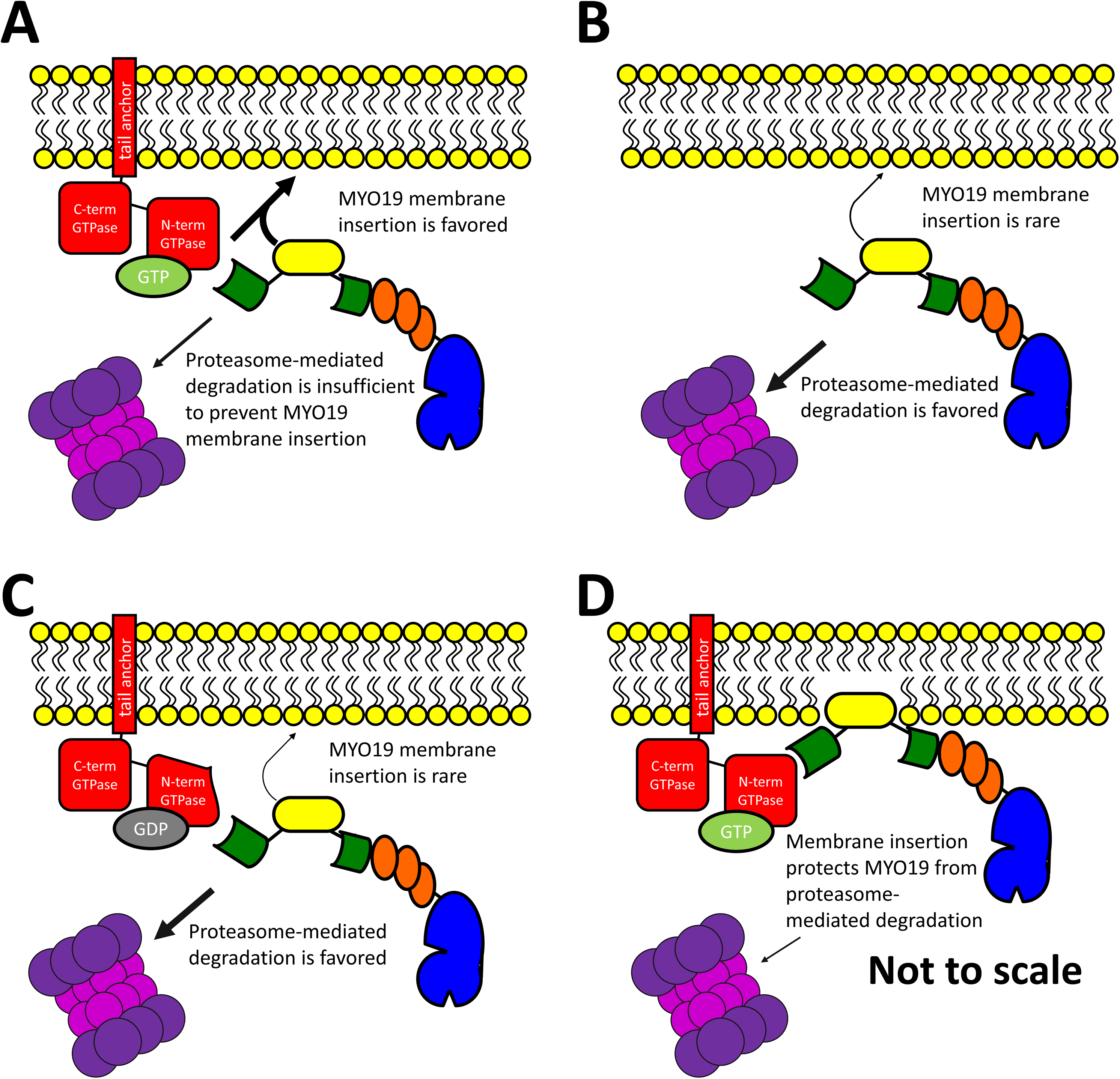
A conceptual model for how Miro/MYO19 interactions facilitate membrane insertion and influence MYO19 degradation. The presence of MYO19 on the mitochondrial outer membrane is a consequence of the balance of membrane-insertion events, and degradation pathways. **A)** Mechanisms that facilitate the removal of MYO19 protein from cytosolic populations to membrane-bound populations, such as interactions with GTP-bound Miro proteins, would result in the presence of MYO19 in the mitochondrial outer membrane. Loss of Miro interactions via **B)** loss of Miro protein or **C)** having Miro proteins in the GDP-bound state would favor proteasome-mediated MYO19 degradation, resulting in decreased mitochondrial MYO19 as well as decreased cellular levels of MYO19 protein. It is also possible that **D)** membrane-inserted MYO19 interacts with Miro in such a way as to protect MYO19 from degradation. These conceptual models are not drawn to scale.

These data all suggest that regulation of the MYO19/Miro interaction would determine how much MYO19 was present on the mitochondrial outer membrane. GTP-bound Miro proteins would enhance mitochondrial localization of MYO19 (Figure 5A), while GDP-bound Miro would shift MYO19 protein towards proteasome-mediated degradation (Figure 5C). If Miro followed the same model of other GTPases, a slow intrinsic Miro GTPase activity would be stimulated by GTPase-activating proteins (GAPs) and nucleotide exchange would be stimulated by proteins with guanine nucleotide exchange factor (GEF) activity. An initial report by Fransson and colleagues indicated that ectopically expressed Trak1 and Trak2 interactions with ectopically expressed Miro1 or Miro2 were not dependent on the nucleotide state of the N-terminal Miro GTPase domain [Fransson et al. 2006]. In contrast, MacAskill and colleagues observed that ectopically expressed Miro1^T18N^ (GDP-mimic) recruits ectopically expressed Trak2 more than Miro1^P13V^ (GTP-mimic) recruits Trak2 [MacAskill et al. 2009a]. It is worth noting that differences exist between the cell types and methods used in each study. Our results indicating that specific mutations to MYO19 disrupt Miro2-binding, that Miro2 switch-mutants disrupt MYO19-binding, and that putative nucleotide-state mutants of Miro2 eliminate MYO19 recruitment are all consistent with a system where the MYO19/Miro2 interaction is dependent on the nucleotide state of the N-terminal Miro2 GTPase. These results are in agreement with similar observations made by Oeding and colleagues for inhibition of MYO19-binding to Miro1 T18N [Oeding et al. 2018]. One recent report exists demonstrating weak intrinsic GTPase activity in human Miro GTPases [Peters et al. 2018]. Additionally, Miro1 interacts with the GEF proteins RAP1GDS1 [Ding et al. 2016] and GBF1 [Walch et al. 2018]. Alterations in the presence or activity of either RAP1GDS1 or GBF1 lead to changes in mitochondrial morphology. To date Miro-specific GEF activity has not been identified biochemically, and little evidence exists for proteins with Miro-specific GAP activity.

Although Miro1 and Miro2 share ∼60% sequence homology and their repertoire of binding-partners may be overlapping, differences in the kinetics of those interactions could indicate related-but-divergent roles in mitochondrial functions, particularly with respect to the role of the actin cytoskeleton and microtubule cytoskeleton in mitochondrial dynamics. Lopez-Domenech and colleagues demonstrated such a functional difference. Ectopic expression of the motor adaptor protein, Trak1, had a larger impact on rescuing mitochondrial positioning in Miro1-knockout MEFs compared to Miro2-knockout MEFs. They were also able to demonstrate that Miro2 knockout had a more drastic impact than Miro1 knockout on cellular MYO19 levels and on the localization of MYO19 to mitochondria [Lopez-Domenech et al. 2018]. We have preliminary evidence that such a functional difference may exist for the Miro proteins with respect to MYO19. We were not able to relocalize MYO19^898-970^ constructs to structures containing mislocalized Miro2 (or vice versa), while Oeding and colleagues were able to recruit MYO19^898-970^ constructs to mislocalized Miro1 [Oeding et al. 2018]. As Miro1 and Miro2 are differentially expressed within cells and across tissues [Fransson et al. 2003] (Supplemental Figure 5), further investigation of the individual properties of Miro1 and Miro2 will be essential to understanding the roles that Miro proteins and MYO19 play in coordinating cytoskeleton-mediated mitochondrial activities.

## Materials and Methods

### Construct generation

MYO19^853-935^-GFP was previously reported [Hawthorne et al. 2016]. Other constructs used in these studies were generated using PFU Ultra II either for PCR mutagenesis or megaprimer PCR insertion [Geiser et al. 2001]. The PCR primers, intended modifications, insert templates, and destination plasmids are listed in Supplemental Table 1 and a schematic of the constructs used can be found in Supplemental Figure 1. pLKO.1-puro-CMV-TagRFP was purchased from Sigma. PEGFP-C1 was purchased from Clontech. GFP-MYO19^824-970^ was previously reported [Quintero et al. 2009]. GFP-Cyto*b*5R was a gift from Nica Borgese [Borgese et al. 2001]. myc-BioID2-MCS was a gift from Kyle Roux [Kim et al. 2016] (Addgene plasmid # 74223; http://n2t.net/addgene:74223; RRID:Addgene_74223). mCherry-LaminA-C-18 was a gift from Michael Davidson (Addgene plasmid # 55068; http://n2t.net/addgene:55068; RRID:Addgene_55068). pRK5-myc-Miro2 (Addgene plasmid # 47891; http://n2t.net/addgene:47891; RRID:Addgene_47891), pRK5-myc-Miro2 T18N (Addgene plasmid # 47897; http://n2t.net/addgene:47897; RRID:Addgene_47897), and pRK5-myc-Miro2 Δ593-618 (Addgene plasmid # 47901; http://n2t.net/addgene:47901; RRID:Addgene_47901) were gifts from Pontus Aspenström [Fransson et al. 2003; Fransson et al. 2006]. EGFP-nesprin-2β was a gift from Gant Luxton. pcDNA3.1-MYO10-HMM-Nanotrap was a gift from Thomas Friedman (Addgene plasmid # 87255; http://n2t.net/addgene:87255; RRID:Addgene_87255) [Bird et al. 2017]. pGEX6P1-mCherry-Nanobody was a gift from Kazuhisa Nakayama (Addgene plasmid # 70696; http://n2t.net/addgene:70696; RRID:Addgene_70696) [Fridy et al. 2014; Katoh et al. 2016]. All generated plasmids were sequenced completely across their coding region.

### Cell culture and transfection

HeLa cells [Scherer et al. 1953] were grown in DMEM high glucose (ThermoFisher) supplemented with 10% fetal bovine serum (Gemini Bio-products) 50 units/mL penicillin, and 50μg/mL streptomycin. Cells were maintained in a humidified incubator at 37°C and 5%CO2. Cells were passaged using 0.25% trypsin-EDTA. For imaging experiments, HeLa cells were grown on 22mm glass coverslips (#1.5) pre-treated with 0.01% poly-L-lysine (Electron Microscopy Sciences) and then coated with 10μg/ml laminin (Cultrex) in PBS for 1h.

Cells were transfected with Lipofectamine 2000 (ThermoFisher) using a modified manufacturer′s protocol. For fixed-cell experiments, 0.1μg of GFP-tagged construct DNA were mixed with 0.2μg of mchr-tagged construct DNA and diluted in Optimem (ThermoFisher) without serum or antibiotics, in a final volume of 150μL per coverslip. In filopodial tip-localization experiments, an additional 0.5 μg of MYO10-Nanotrap DNA was added. For live-cell experiments (FRAP or PARF), 0.3μg of GFP-tagged construct DNA were mixed with 0.7μg of mchr-tagged construct DNA and diluted in Optimem (ThermoFisher) without serum or antibiotics, in a final volume of 150μL. In all experiments, 4μL of Lipofectamine was diluted into 150μL of Optimem without serum or antibiotics and then mixed with the DNA dilution. Complexes were allowed to form for 5 minutes at room temperature. The entirety of the DNA/reagent mix was added drop-wise to a well of a 6-well plate. Cells were used for experimentation 18 to 30 hours after transfection.

### Establishment of cell lines stably expressing MYO19^824-970^-BioID2

Lentiviral particles were generated by the Duke University Viral Vector Core Facility as previously described [Vijayraghavan and Kantor 2017]. HeLa cells were grown in growth media supplemented with 8μg/ml hexadimethrine bromide, and infected at a 10:1 multiplicity of infection. After two days of infection, cells were grown in growth media supplemented with 2μg/ml puromycin for 2 weeks. Stable expression of MYO19^824-970^-BioID2 was verified via western blotting and immunostaining. Cells were then trypsinized, resuspended in growth media supplemented with 10%DMSO and 40% fetal bovine serum, aliquoted, and frozen.

### Immunostaining

MYO19^824-970^-BioID2 stable cells were plated on coverslips as previously described, and grown in the presence of 50μM biotin overnight. Prior to fixation, cells were stained with 50nM Mitotracker CMXRos diluted in growth media for 15 minutes, and then incubated in growth media for 15 minutes. Cells were then quickly washed with 37°C PBS, fixed in 4% paraformaldehyde in PBS for 10 minutes at 37°C, and permeabilized in 0.5% TritonX-100 in PBS for 10 minutes. Cells were then blocked (PBS, 1% bovine serum albumin, 0.2% TritonX-100) for 30 minutes, and incubated in mouse α-myc primary antibody (1:250 in blocking buffer, Invitrogen R950-25) for 1 hour. Cells were then washed in PBS with 0.2% TritonX-100 four times for five minutes, followed by a 1 hour incubation with ALEXA350-streptavidin (1:250, ThermoFisher), ALEXA488 goat α-mouse secondary antibody (0.2μg/ml, JacksonImmuno), and ALEXA647 phalloidin (1:1000, ThermoFisher) diluted in blocking buffer. After four 5-minute washes in PBS with 0.2% TritonX-100, the cells were mounted on coverslips using PBS in 80% glycerol and 0.5% N-propyl gallate. Coverslips were sealed onto slides with Sally Hansen Tough as Nails clear nail polish.

HeLa cells expressing MYO19-GFP and/or mchr-Miro2 constructs were assayed for endogenous MYO19 expression using a rabbit α-human MYO19 antibody (0.9μg/ml, abcam #174286). The day after transient transfection, cells were quickly washed with 37°C PBS, fixed in 4% paraformaldehyde in PBS for 10 minutes at 37°C, and permeabilized in 0.5% TritonX-100 in PBS for 10 minutes. Cells were then blocked (PBS, 5% donkey serum) for 1 hour, and the incubated in rabbit α-MYO19 primary antibody (1:250 in blocking buffer) for 90 minutes. Cells were then washed in PBS four times for five minutes, followed by a 1-hour incubation with ALEXA647-donkey α-rabbit secondary antibody (0.5μg/mL) and DAPI (60nM) diluted in blocking buffer. After four 5-minute washes in PBS, the cells were mounted on coverslips using PBS in 80% glycerol and 0.5% N-propyl gallate. Coverslips were sealed onto slides with Sally Hansen Tough as Nails clear nail polish.

### Streptavidin pulldown

MYO19^824-970^-BioID2 stable cells were plated in 150mm cell culture dishes. Once they reached approximately 80% confluency, they were grown in the presence of 50μM biotin overnight (minimum of 8 hours). Biotinylated proteins were purified as previously described [Mehus et al. 2016]. Briefly, cells were lifted from the plate using 0.25% trypsin EDTA, and washed with PBS via centrifugation (700xg, 4°C, 10 minutes). Cells were resuspended in lysis buffer (50mM Tris HCl, 500mM NaCl, 0.2% SDS, 1mM DTT, 10 μg/ml aprotinin, 10μg/ml leupeptin, 1mM PMSF, pH 7.4). TritonX-100 was added to a final concentration of 2%, and the lysate was mixed by inversion. Samples were then placed on ice and sonicated twice (Branson Sonifer 450 Digital, 40% amplitude, 30% duty cycle, 30, 0.5s pulses), with a 2-minute incubation on ice between sonications. Lysates were diluted ∼4-fold with 50mM Tris HCl, pH7.4 prior to a third round of sonication. Insoluble material was removed from the lysate via centrifugation (16,5000g, 4°C, 10 minutes).

Magnetic streptavidin beads were washed in 50mM Tris HCl, pH7.4 prior to incubation with the lysate supernatant at 4°C with gentle rocking overnight. The sample were then collected to the side of the tube using a magnetic stand. The beads were washed (8-minute incubation time) twice with wash buffer 1 (2% SDS), once with wash buffer 2 (50mM Hepes, 500mM NaCl, 1mM EDTA, 1% TritonX-100, 0.1% deoxycholic acid, pH 7.5), and once with wash buffer 3 (10mM Tris HCl, 250mM LiCl, 1mM EDTA, 0.5% NP-40, 0.5% deoxycholic acid, pH 7.4). The beads were then resuspended in 50mM Tris HCl, pH7.4, with 10% saved for western blot analysis. The remaining sample was resuspended in 50mM ammonium bicarbonate in preparation for mass spectrometry analysis.

### Mass spectrometry sample preparation, analysis, data filtering, and bioinformatics

Samples were subjected to in-solution Trypsin digestion following a RapiGest SF Surfactant (Waters Corporation, Part# 186001861) elution. Trypsinzation was achieved by incubating eluted protein with excess trypsin (Promega V5111) at 37°C overnight (16-18 hours). Subsequently, eluted peptides were desalted using a Pierce C-18 spin column (Thermo 89870) and followed by an ethyl acetate cleanup step to remove detergents. Peptides were dried down and then re-suspended in 2% Formic Acid LC-MS grade water solution for mass spec analysis.

Peptides were separated using reverse-phase nano-HPLC by a nanoACQUITY UPLC system (Waters Corporation). Peptides were trapped on a 2 cm column (Pepmap 100, 3μM particle size, 100 Å pore size), and separated on a 25cm EASYspray analytical column (75μM ID, 2.0μm C18 particle size, 100 Å pore size) at 45°C. The mobile phases were 0.1% formic acid in water (buffer A) and 0.1% formic acid in acetonitrile (buffer B). A 180-minute gradient of 2-25% buffer B was used with a flow rate of 300nl/min. Mass spectral analysis was performed by an Orbitrap Fusion Lumos mass spectrometer (Thermo Scientific). The ion source was operated at 2.4kV and the ion transfer tube was set to 300°C. Full MS scans (350-2000 m/z) were analyzed in the Orbitrap at a resolution of 120,000 and 1e6 AGC target. The MS2 spectra were collected using a 1.6 m/z isolation width and were analyzed either by the Orbitrap or the linear ion trap depending on peak charge and intensity using a 3s TopSpeed CHOPIN method [Davis et al. 2017]. Orbitrap MS2 scans were acquired at 7500 resolution, with a 5e4 AGC, and 22ms maximum injection time after HCD fragmentation with a normalized energy of 30%. Rapid linear ion trap MS2 scans were acquired using an 4e3 AGC, 250ms maximum injection time after CID 30 fragmentation. Precursor ions were chosen based on intensity thresholds (>1e3) from the full scan as well as on charge states (2-7) with a 30-s dynamic exclusion window. Polysiloxane 371.10124 was used as the lock mass.

Raw mass spectrometry data were searched against the Swiss-Prot human sequence database (released 2/2017) using MaxQuant version 1.6.2.3. The parameters for the search were as follows: specific tryptic digestion with up to two missed cleavages, static carbamidomethyl cysteine modification, variable protein N-terminal acetylation and methionine oxidation, Label Free Quantification (LFQ) and match between runs were enabled. Protein identifications were filtered for a false discovery rate (FDR) of 1%, and potential contaminants and decoys were removed. To score candidate protein-protein interactions, SAINTq version 0.0.4 using LFQ values was used and then filtered for a 10% FDR.

The mass spectrometry proteomics data have been deposited to the ProteomeXchange Consortium via the PRIDE partner repository with the dataset identifier XXXXXXX. ***Will be completed to coincide with publication.***

### Western blot analysis

5x Laemelli sample buffer (310mM Tris-HCl pH 6.8, 10% SDS, 50% Glycerol, 2.5% Bromophenol Blue, 7.5% β-Mercaptoethanol) was added to all cell pellet and pulldown samples prior to boiling at 95°C for 5 minutes. Samples were loaded onto a 4-12% Bis-Tris NuPAGE gels, transferred to nitrocellulose, and transfer was verified by Ponceau staining (0.5% Ponceau Red-S, 2% Acetic acid in H2O). Membranes were incubated in blocking buffer (PBS, 0.2% TritonX-100, 1% bovine serum albumin, pH 7.4) for 20 minutes, incubated with streptavidin-HRP (1:5000, ThermoFisher) for 40 minutes, and then washed in PBS three times for 5 minutes. Blots were then exposed to Clarity ECL (Bio-Rad) for 5 minutes, and visualized using the BioRad ChemiDoc MP imager.

### Fixed-cell preparation and image analysis

Transfected cells grown on coverslips were stained with 100nM Mitotracker DeepRed FM diluted in growth media for 10 minutes, and then washed in growth media for 10 minutes. Cells were then washed once with PBS quickly prior to fixation in PBS with 4% paraformaldehyde 37°C. After permeabilization in PBS with 0.5% TritonX-100 for 5 minutes, cells were stained in PBS with 75nM DAPI for 10 minutes, washed in PBS four times for five minutes, and mounted on slides using PBS in 80% glycerol and 0.5% N-propyl gallate. Coverslips were sealed onto slides with Sally Hansen Tough as Nails clear nail polish. For experiments using MYO10-Nanotrap constructs, cells were not stained with Mitotracker DeepRed FM, but instead were stained with 6.6nM ALEXA647 phalloidin during the DAPI staining step.

Images were acquired using an Olympus IX-83 microscope and a PLAN APON 60x/1.42NA DIC objective or a PLAN FLUOR 40x/1.3 NA objective. Cells were illuminated using an EXFO mixed gas light source, Sedat Quad filters (Chroma), and Sutter filter wheels & shutters. Images were acquired using a Hamamatsu ORCA-Flash 4.0 V2 sCMOS camera, and Metamorph imaging software to control the system components. In all instances, exposure times maintained constant across experimental data sets.

### Quantitative image analysis of fixed cells

The ratio of organelle-localized fluorescence to cytoplasm-localized fluorescence (mito/cyto ratio) was calculated [Hawthorne et al. 2016] by measuring the integrated density of in-focus organelles and cytosol with 3×3 pixel boxes using FIJI [Schindelin et al. 2012]. For each cell measurement, the ratio was calculated from five separate regions within one cell and then averaged. A ratio with a value greater than 1 indicates that the signal is more concentrated in the organelle region than in the cytosol region. The same analysis method was used to calculate the filopodial tip to cytosol ratio.

The fraction of cells within a population showing a cytoplasmic, mitochondrial, or mixed pattern of GFP fluorescence was calculated by manually scoring 60-100 cells/coverslip for each phenotype. The data collector was blinded to the identity of the samples during data collection. The percentage of cells in each sub-population was calculated by dividing the number of cells with a particular phenotype by the total number of cells counted for that coverslip.

The fluorescence intensity of ALEXA350-streptavidin labelling was calculated by selecting perinuclear mitochondrial regions in Metamorph and calculating the average pixel fluorescence in those regions. Control HeLa and MYO19-BioID2 HeLa cells were stained in parallel under identical staining conditions, and all images used in these analyses were captured under identical imaging conditions.

The relative fluorescence along a line crossing the nuclear envelope was calculated using the “Plot Profile” tool in FIJI. The relative brightness at each point along the line was calculated by subtracting the minimum brightness along the line from the brightness at that point. Then that background-adjusted brightness was divided by the background-adjusted maximum fluorescence level along the line.

### Live-cell imaging & kinetic analysis

Transfected HeLa cells were enclosed in a Rose chamber [Rose et al. 1958] filled with Optimem without phenol red containing 50 units/mL penicillin, and 50μg/mL streptomycin. Images were collected on an Olympus FV1200 laser scanning confocal microscope outfitted with a PLAN APON 60x/1.4NA objective at a frame-rate of 1 frame every 2 seconds, or 1 frame every 10 seconds (depending on the sample). After the first 30 frames, regions of interest were illuminated for photobleaching at high laser power for 1 second. FIJI was used to identify regions of interest and calculate average fluorescence intensity for each time point. Relative fluorescence at each time point was measured by determining the fluorescence intensity relative to the frame prior to photobleaching (t = 0s). Data were corrected for photofading due to imaging [Applewhite et al. 2007] and averaged together. The mean gray value relative to t = 0 was plotted over time and fit to the function y = a*(1 - *e*^−bt^) + c*(1 - *e*^−dt^) +e. The half-life (t1/2) was calculated using from the exponential rise function by calculating the value of t when y = 0.5. Immobile fraction was calculated by using the exponential fit of the data to determine the value of the asymptote being approached.

PARF analysis of transfected HeLa cells was completed as previously described [Singh et al. 2016]. Cells were mounted in an open-top Rose chamber filled with with 300μL KHM buffer (pH 7.4, 110 mM potassium acetate, 20 mM HEPES, 2 mM MgCl2). Images were acquired at the rate of one frame every 2 seconds. After the first 20 frames, digitonin was added (t=0) to a final concentration of 25μM. FIJI was used to select regions of cells not including the nuclei where average fluorescence intensity was measured for each frame. Fluorescence intensity relative to the frame prior to permeabilization (t = 0) was calculated for multiple experiments. These data were corrected for photofading due to imaging, and averaged together. The mean gray value relative to t = 0 was plotted over time and fit to a double exponential function of the form y = a + b\**e*^(−ct)^ + d\**e*^(−ft)^. The half-life (t1/2) was calculated using the exponential decay function calculating the value of t when y = 0.5. Immobile fraction was calculated by using the exponential fit of the data to determine the value of the asymptote being approached.

### Sequence alignment

The MYO19 amino acid sequences from seven vertebrates including mammals (human: Q96H55, mouse: NP_079690, dog: XP_022278968, cow: NP_001019672), chicken (XP_024997806.1), *Xenopus laevis* (AAH92309.1), and zebrafish (XP_021332051.1), were manually trimmed to the C-terminal sequence containing the third IQ motif and containing the MyMOMA domain. Sequences were aligned using Clustal Omega with standard settings [Sievers et al. 2011]. Amino acid sequences for the N-terminal region of Miro2 (human: NP_620124.1, mouse: NP_001351879.1, dingo: XP_025272941, cow: NP_847886, chicken: NP_001074335.1, *Xenopus tropicalis:* NP_001006725.1, and zebrafish: XP_021332051.1) were aligned using Clustal Omega with standard settings.

### Quantification, analysis, and statistics

Metamorph (Universal Imaging) and FIJI were used for image analysis. Data points are expressed as means ± standard error of the mean. Data were compared by Student’s t-test, Tukey analysis, or Dunnett’s analysis using Kaleidagraph. Exponential fits for PARF and FRAP analysis were performed using Kaleidagraph. Parameters calculated from exponential fits are expressed as mean ± the SEM-adjusted fit [Singh et al. 2016], by generating curve fits of the mean + SEM and mean − SEM, and reporting the error as the difference between the value calculated from the mean fit and the value calculated from the SEM-adjusted fit. All images were prepared for publication using FIJI, Metamorph, Photoshop, or some combination of these software packages.

## Supporting information

Bocanegra_Supplemental Table 1

Bocanegra_Supplemental Table 2

## Acknowledgements and author contributions

OAQ was supported by a grant from the National Institute of General Medical Sciences at the NIH (R15 GM119077) and by funding from the University of Richmond School of Arts & Sciences. JLB and JMC were supported by the NIGMS grant to OAQ. BMF was supported by the Robert F. Smart Award from the Biology Department, and funding from the School of Arts & Sciences at the University of Richmond. NRM and ELS were supported by funding from the University of Richmond School of Arts and Sciences. TYT was supported by an HHMI Gilliam Fellowship for Advanced Study. We would like to thank Uri Manor, Anna Hatch, Jenci Hawthorne, Rebecca Adikes, Sarah Rice, Susan Walsh, and Andrew Moore for critical reading and discussions related to this manuscript. Hypotheses and experiments in this study were conceived by OAQ. Experiments were performed by JLB, BMF, NRM, JMC, ELS, TYT, and OAQ. Data analysis and manuscript preparation were completed by JLB, BMF, NRM, JMC, TYT, MBM, and OAQ. Except for the proteomics analysis, all experiments for the initial submission were completed during the 10-week 2018 summer research session at University of Richmond. We would also like to that Edward Salmon for his example and inspiration.

---

**Supplemental Figure 1.**
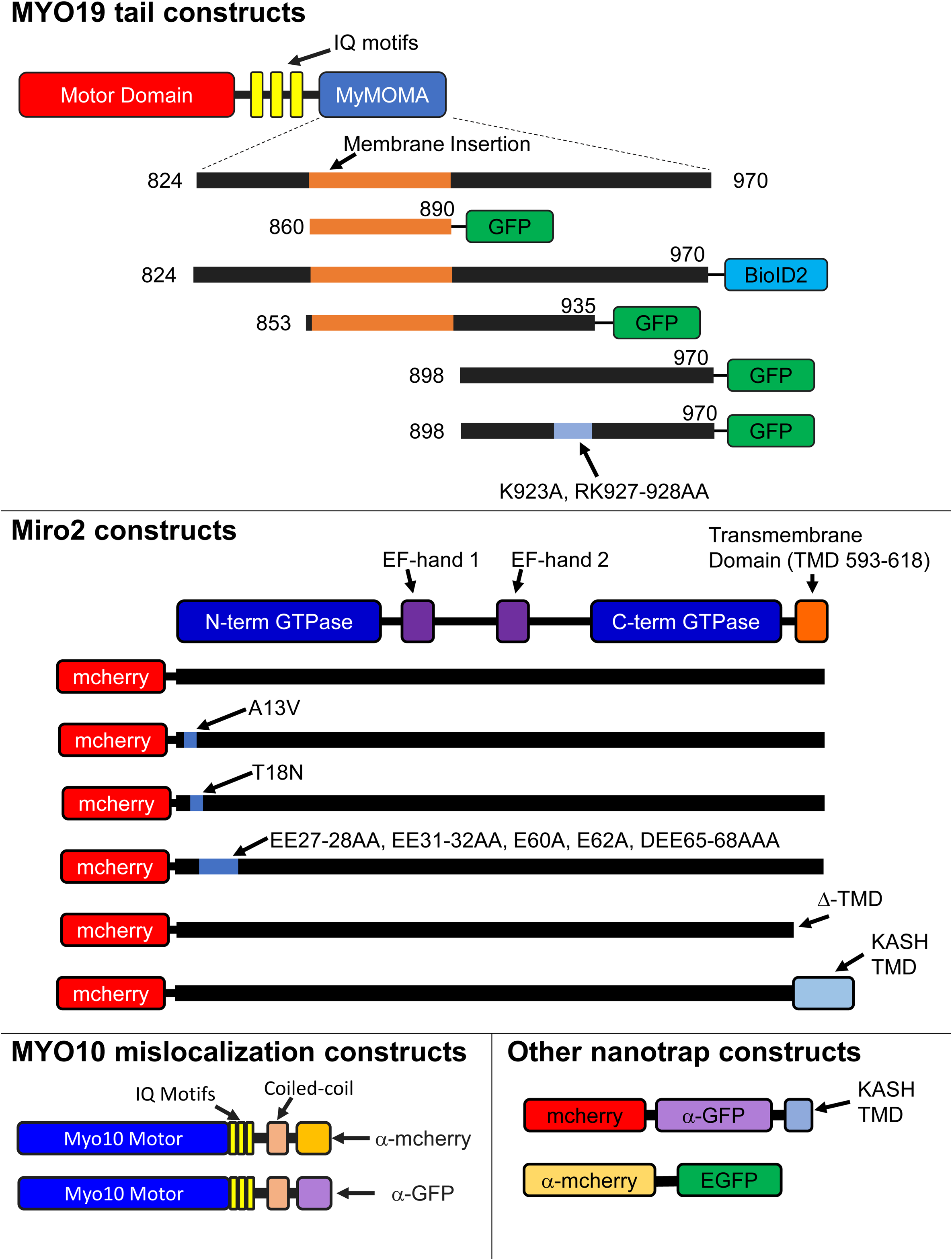
Schematic diagrams of constructs used in these studies. Multiple epitope-tagged expression constructs were generated for these studies using the plasmids and primer sets listed in Supplemental Table 1, as described in the Materials and Methods section.

**Supplemental Figure 2.**
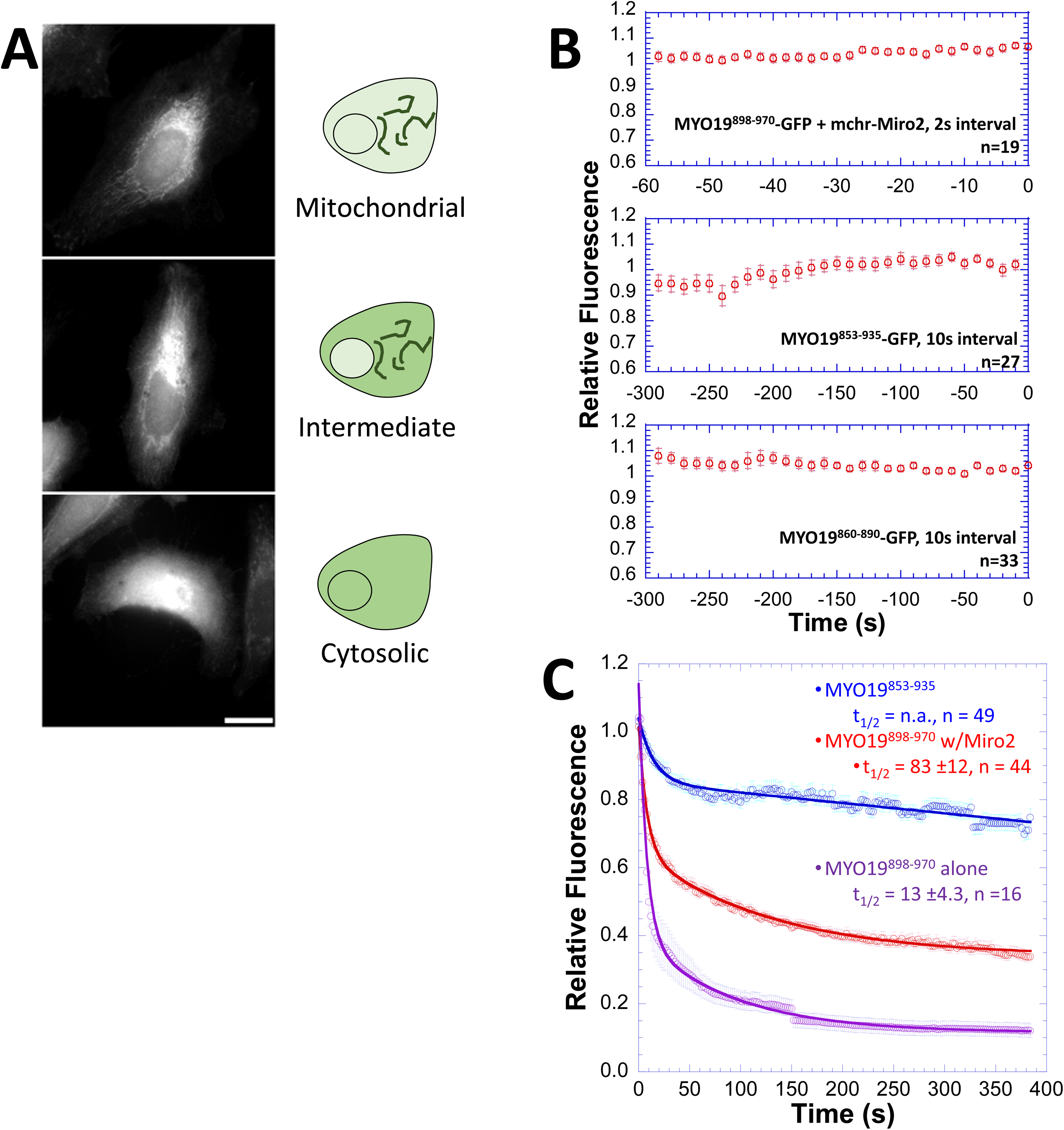
Controls for MYO19 image analysis. **A)** Representative images for the mitochondrial localization categories used in the population analysis of MYO19 localization. Localization of MYO19 to the mitochondrial outer membrane was classified into three levels: mitochondrial, intermediate, and cytosolic. The mitochondrial pattern displayed little cytosolic labeling while the cytosolic pattern displayed almost no mitochondrial labeling distinguishable from the cytosolic signal. The intermediate cells exhibited localizations patterns somewhere in between the two extremes—identifiable mitochondrial labeling with substantial cytosolic signal as well. Scale bar, 20 μm. **B)** Prebleach fluorescence intensity curves for MYO19^853-935^-GFP, MYO19^860-890^-GFP, and MYO19^8998-970^-GFP coexpressed with mchr Miro2. The curves represent the photofading correction of the 30 frames prior to photobleaching. Note that time intervals are different between MYO19^898-970^-GFP with mchr-Miro2 (2s interval) compared to the other two constructs (10s interval). Error bars represent standard error of the mean. **C)** PARF analysis shows that MYO19^853-935^-GFP dissociates more slowly and has a larger immobile fraction than either MYO19^898-970^-GFP or MYO19^898-970^-GFP coexpressed with mchr-Miro2. Error bars represent standard error of the mean. PARF kinetic parameters are listed in Table 2.

**Supplemental Figure 3:**
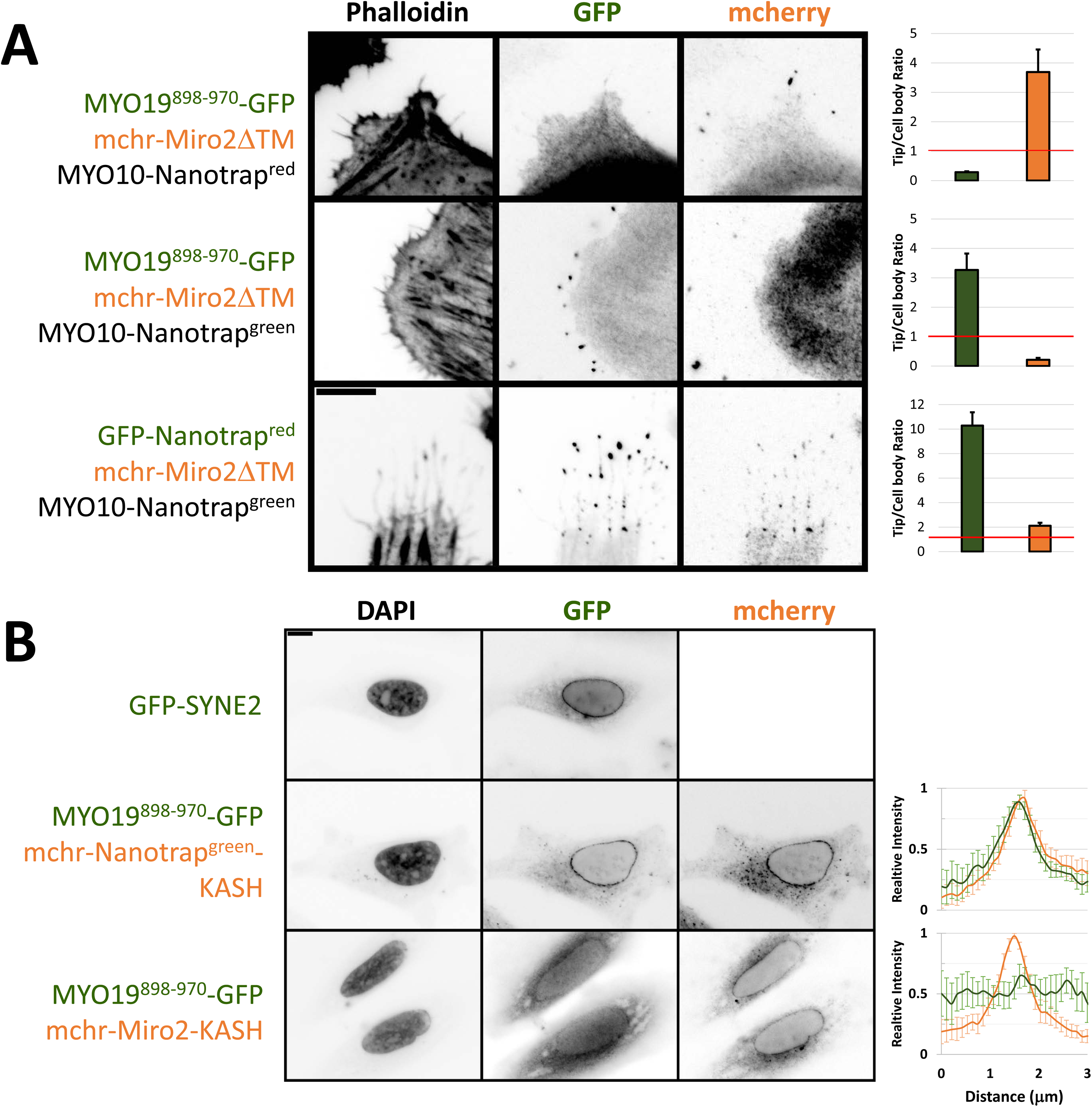
Mislocalization of Miro2 to either filopodial tips or the nuclear envelope does not recruit MYO19^898-970^-GFP to either location. **A)** mchr-Miro2 was mislocalized by removing the C-terminal mitochondrial outer membrane domain (mchr-Miro2ΔTM). When coexpressed with MYO10-Nanotrap^red^ and MYO19^898-970^-GFP, mchr-Miro2ΔTM localized to filopodial tips but GFP signal did not concentrate with it. When MYO19^898-970^-GFP, mchr-Miro2ΔTM, and MYO10-Nanotrap^green^ were coexpressed, GFP signal collected at filopodial tips but mcherry signal did not. When GFP-Nanotrap^red^, mchr-Miro2ΔTM, and MYO10-Nanotrap^green^ were coexpressed, both GFP and mcherry signal localized to filopodial tips. Bar graphs indicate the ratio of filopodial tip fluorescence to cell body fluorescence for 6 filopodia in the image. Ratios above 1 indicate fluorescence concentration at the filopodial tip. Note that the only instance where the TCB ratio is above one in both channels is GFP-Nanotrap^red^/mchr-Miro2ΔTM/MYO10-Nanotrap^green^. Error bars represent standard error of the mean. B) GFP-nesprin-2β contains a C-terminal KASH domain which localizes the protein to the cytosolic face of the nuclear membrane. Cells coexpressing mchr-Nanotrap^green^-KASH and MYO19^898-970^-GFP show mcherry localization and GFP localization at the nuclear membrane. Cells coexpressing mchr-Miro2-KASH and MYO19^898-970^-GFP show mcherry localization at the nuclear membrane but not concentration of GFP signal. Linescans across the border between the nucleus and cytosol indicate regions of high brightness at the nuclear periphery for both channels in MYO19^898-970^-GFP/mchr-Nanotrap^grenn^-KASH cells, but not in MYO19^898-970^/mchr-Miro2-KASH cells. Graphs represent the average relative brightness across a 3μm region for six cells. Scale bar = 10 µm.

**Supplemental Figure 4:**
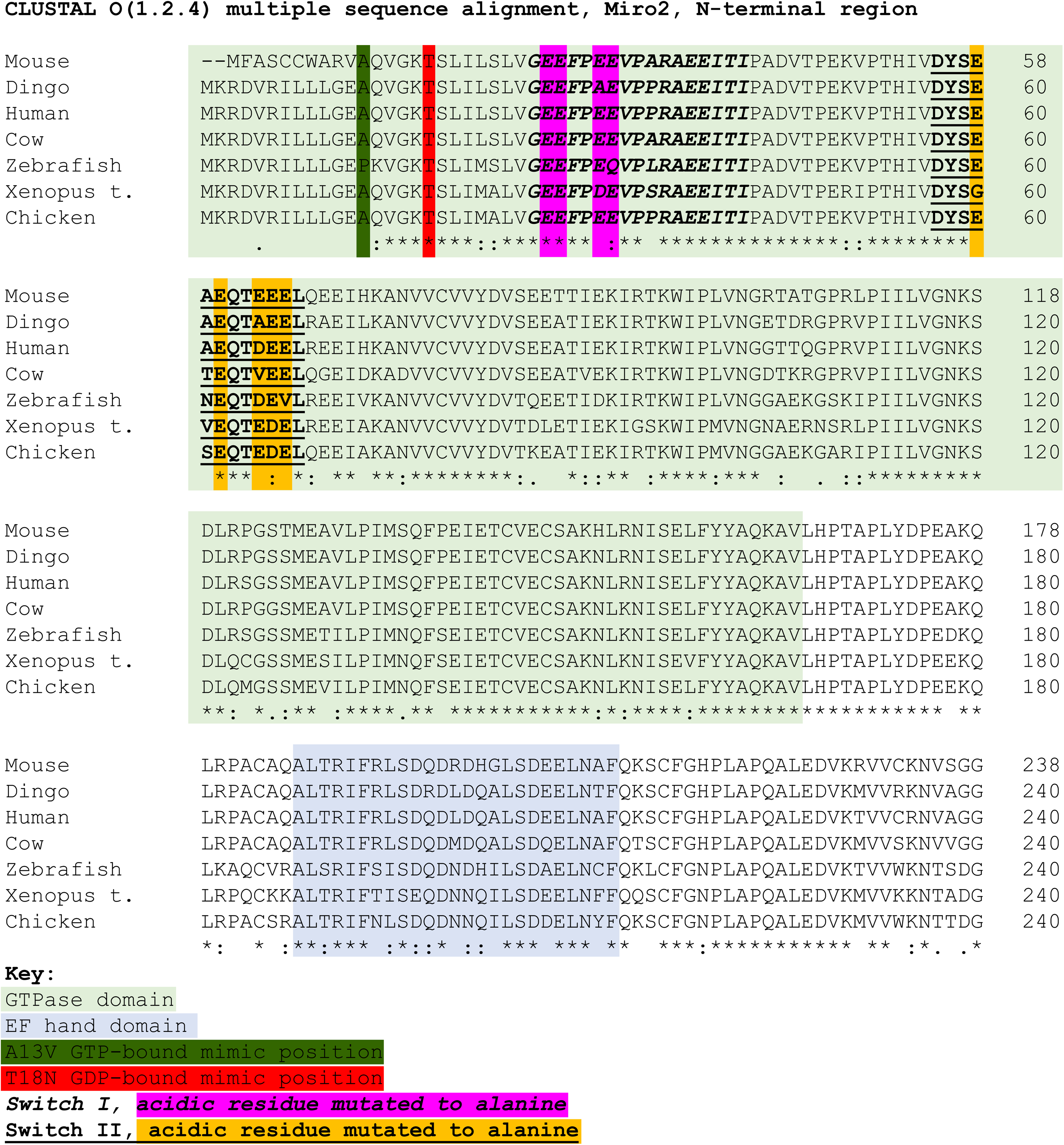
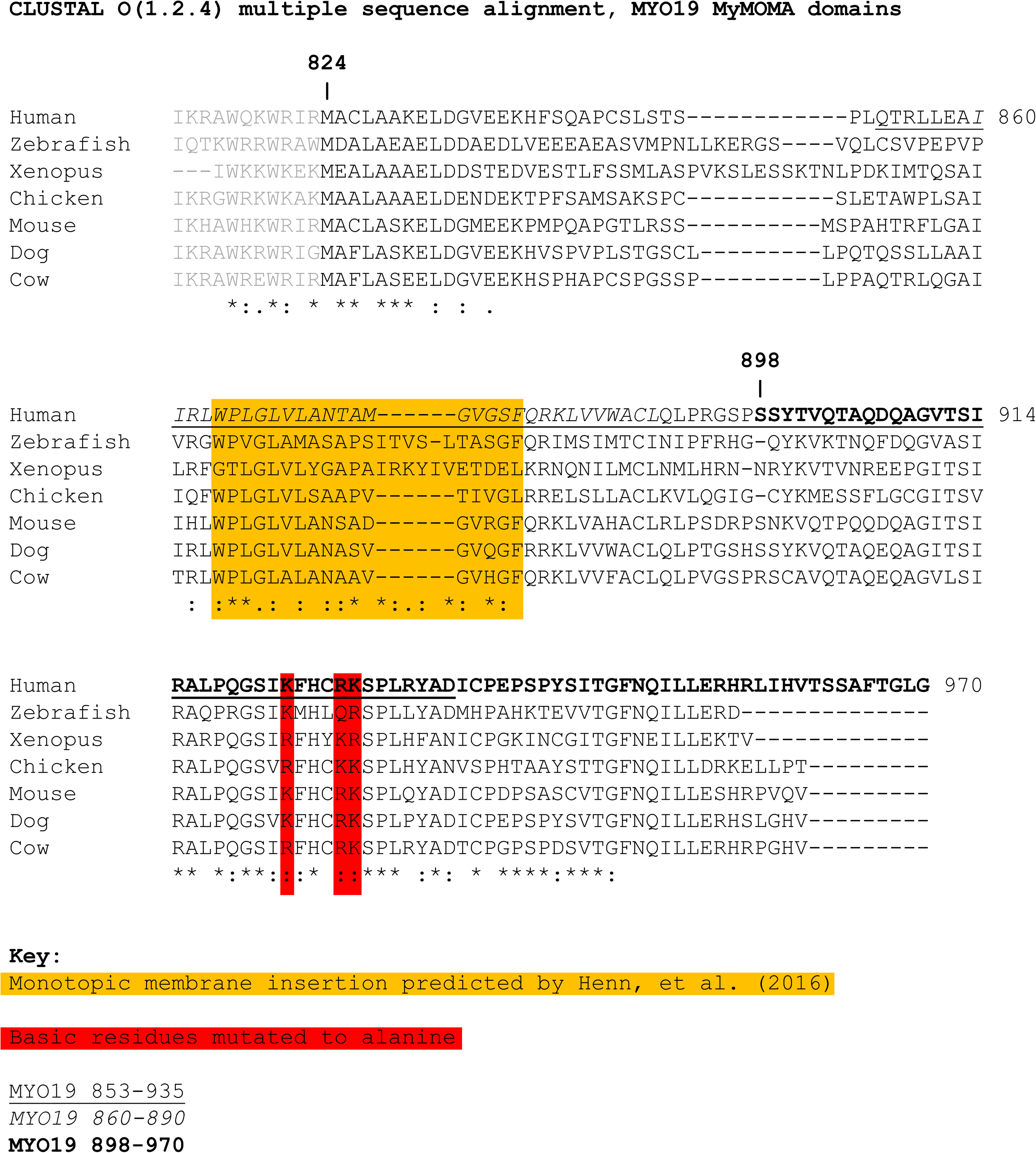
Multiple sequence alignment of Miro2 and MYO19. **A)** Seven vertebrate Miro2 N-terminal GTPase domains reveals well-conserved acidic residues within switch I and switch II. Full-length Miro2 sequences for seven species (human: NP_620124.1, mouse: NP_001351879.1, dingo: XP_025272941, cow: NP_847886, chicken: NP_001074335.1, *Xenopus tropicalis:* NP_001006725.1, and zebrafish: XP_021332051.1) were aligned using Clustal Omega with standard settings. **B)** The MYO19 amino acid sequences from seven vertebrates (human: Q96H55, mouse: NP_079690, dog: XP_022278968, cow: NP_001019672, chicken: XP_024997806.1, *Xenopus laevis:* AAH92309.1, and zebrafish: XP_021332051.1) were manually trimmed to the C-terminal sequence containing the third IQ motif and containing the MyMOMA domain. Sequences were aligned using Clustal Omega with standard settings. Positions with a single, fully conserved residue are indicated by an asterisk (*). Positions with strongly conserved residues are indicated by a colon (:). Positions with weakly conserved residues are indicated by a period(.).

**Supplemental Figure 5:**
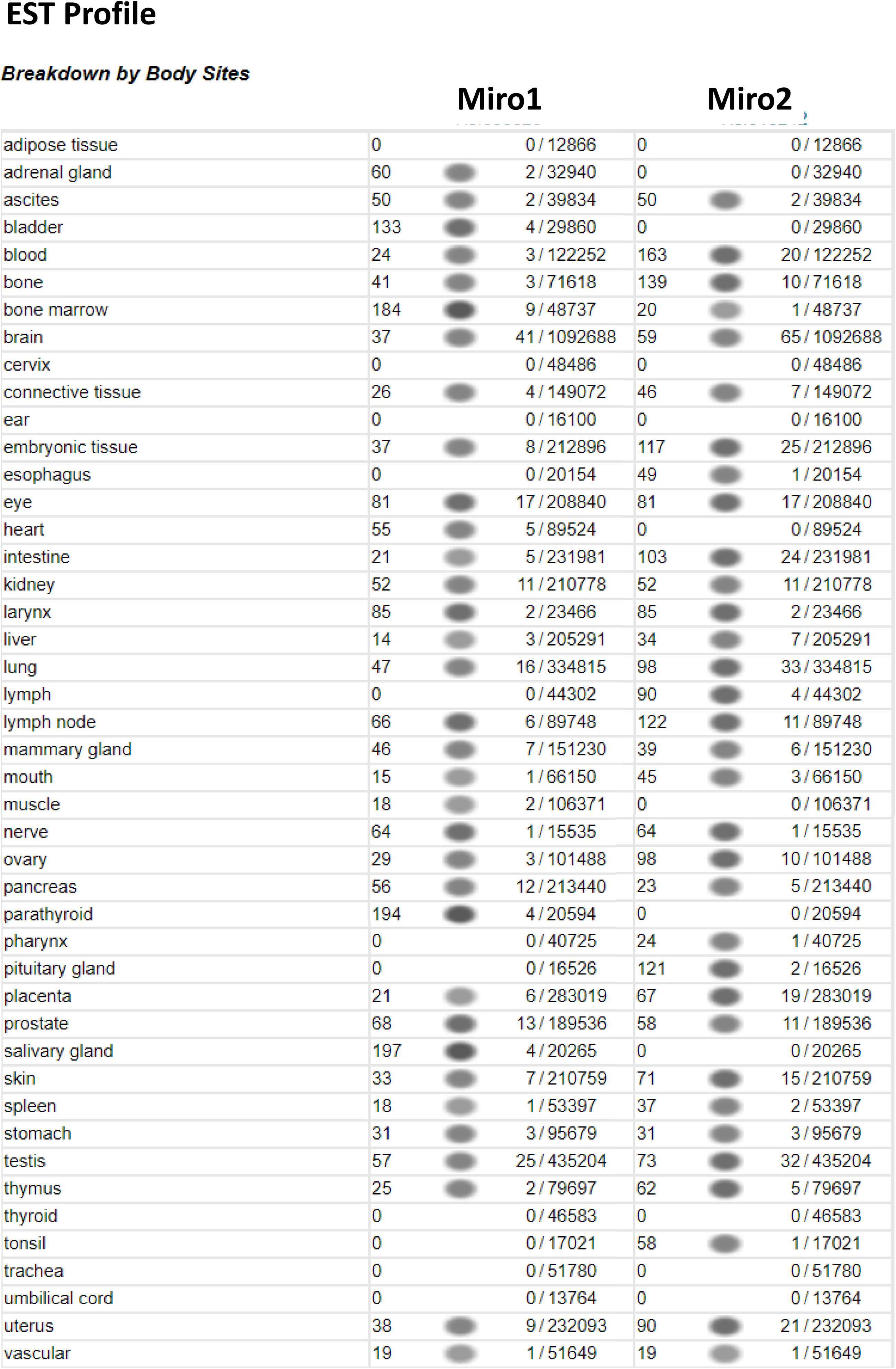
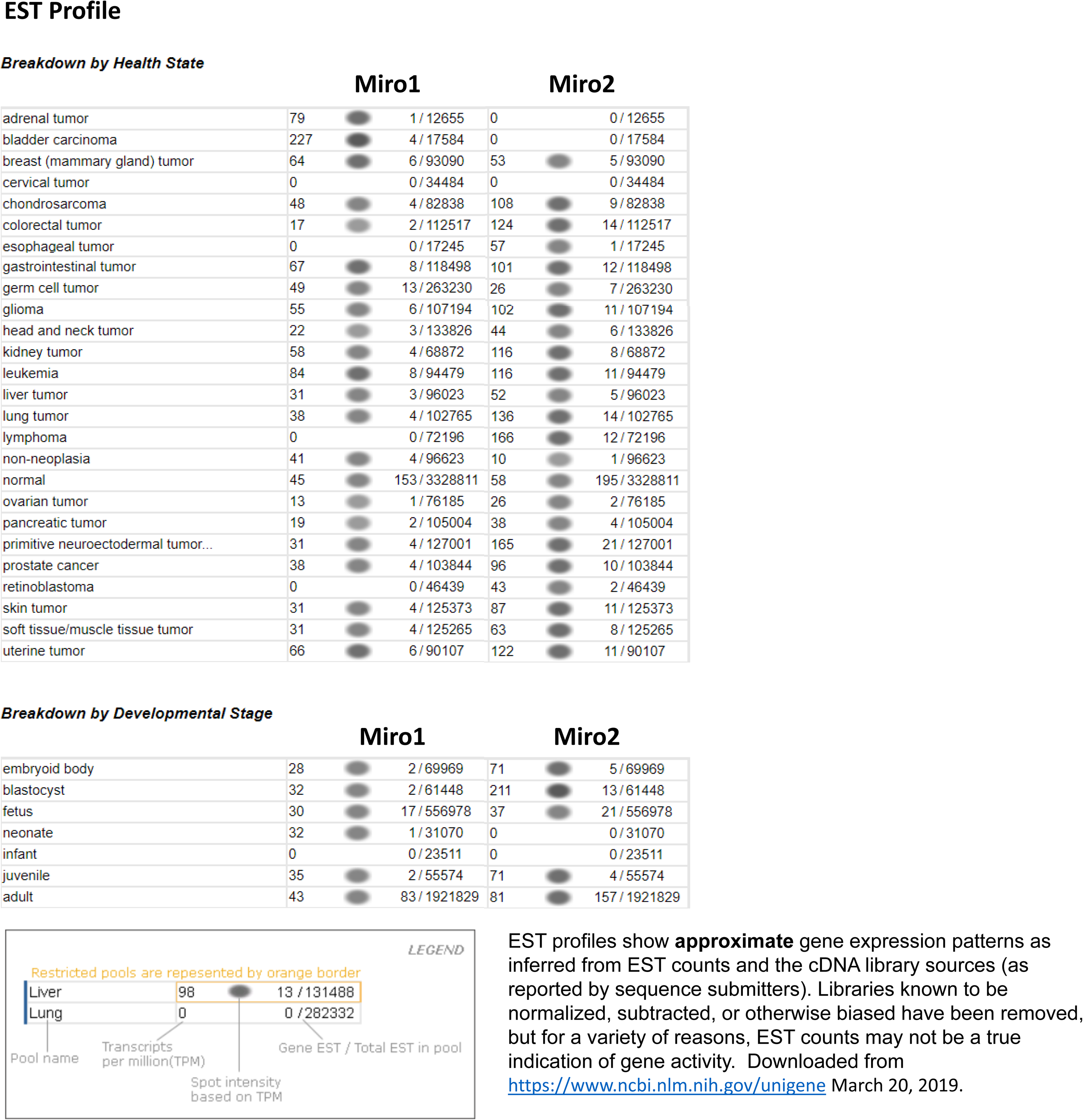
Expression analysis based on expressed sequence tags (EST) suggest differential expression across multiple tissues and cellular states between Miro1 and Miro2. Human gene expression profiles by Unigene estimate relative gene expression levels for Miro1 (Hs.655325) and Miro2 (Hs.513242). These estimates suggest differences in expression levels between Miro1 and Miro2 in A) a variety of tissues as well as B) by health state and developmental stage. Results downloaded from https://www.ncbi.nlm.nih.gov/unigene on March 30, 2019.

**Supplemental Table 1: Primers and plasmids used to generate the expression constructs for these studies.**

**Supplemental Table 2: Promiscuous biotinylation results for MYO19^824-970^-BioID2 (Excel file).** HeLa cells stably expressing MYO19^824-970^-BioID2 were grown in the presence of 50µM biotin overnight. The fraction of biotinylated proteins were purified, analyzed by mass spectrometry, and compared to biotin-exposed cells not expressing a BioID2 construct. This excel file contains the MaxQuant and SAINT analysis results, as well as THEBIOGRID affinity pulldown data for MYO19 (accessed on March 29, 2019), which was used as a comparison in order to identify putative MYO19 interactors. Proteins identified by Oeding et al. [Oeding et al. 2018] are indicated in bold on the “SAINT analysis” tab.

