## Supplementary material for "The MyMOMA domain of MYO19 encodes for distinct Miro-dependent and Miro-independent mechanisms of interaction with mitochondrial membranes": Bocanegra_Supplemental Table 1

**Supplemental Table 1: Primers and plasmids used to generate the expression constructs for these studies**

| Primer Type | Insert plasmid | Destination plasmid | Modification to Destination plasmid | Final Plasmid | Primer Sequence |
|---|---|---|---|---|---|
| Megaprimer | myc-BioID2-MCS (Addgene #74223) | pLKO.1-puro-CMV-TagRFP (Sigma #SHC012) | Replacement of tRFP with myc-BioID2 | URCF1 pLKO.1 myc-BioID2 | AGATCCGCTAGCGACGCCACCATGGAACAAAAACTCATCTCAGAAGAGG<br>ATGGGGCCCCCGTTCTGCAGTCAGCGGTTTAAACTTAAGCTTGGTACCGA |
| Megaprimer | GFP-CytoB5RR (Borgese) | URCF1 pLKO.1 myc-BioID2 | Addition of linker between myc and BioID2<br>SGLRSRAQASNSDLEGGGGSGGGGSGGGGS | URCHb4 pLKO.1 myc-linker-BioID2 | ATGGAACAAAAACTCATCTCAGAAGAGGATCTCTCCGGACTCAGATCTCGAGCTC<br>CTTCAGCCAGATCAGGTTCTTGAAGTCGGATCCGCCACCTCCAGATC |
| Megaprimer | GFP-MYO19$^{824-970}$ | URCHb4 pLKO.1 myc-linker-BioID2 | Insertion of MYO19$^{824-970}$ between linker and BioID2 | URCL5 pLKO.1 myc-linker-MYO19$^{824-970}$-BioID2 | TGGAACAAAAACTCATCTCAGAAGAGGATCTCATGGCCTGCCTTGCTGCTAAAGA<br>CTTGAGCTCGAGATCTGAGTCCGGACCCCAGCCCAGTGAAGGC |
| Deletion | n/a | URO13 MYO19$^{866-970}$-EGFP | Removal of MYO19 amino acids 866-897 | URDS10 MYO19$^{898-970}$-EGFP | CTCGAGCTCAAGCTTATGAGTAGCTACACTGTCCAG<br>CTGGACAGTGTAGCTACTCATAAGCTTGAGCTCGAG |
| Quickchange | n/a | URDS10 MYO19$^{898-970}$-EGFP | Mutation of well-conserved basic residues<br>in the MYO19 Miro-binding sequence | UREM4 MYO19$^{898-970}$-EGFP K923A RK927-28AA | GCTGCCTCAGGGATCGATAGCGTTTCACTGCGCAGCGTCTCCACTGCGGTATGC<br>GCATACCGCAGTGGAGACGCTGCGCAGTGAAACGCTATCGATCCCTGAGGCAGC |
| Megaprimer | mCherry-LaminA-C-18 (Addgene #55068) | pRK5-myc-Miro2 (Addgene #47891) | Addition of mcherry | URDT12 mchr-Miro2 | ATCTCCGAGGAGGACCTGGGATCTATGGTGAGCAAGGGCGAGGAG<br>GATGCGCACGTCCCGCCTCATGCTTCCGCTTCCGCCGG |
| Megaprimer | mCherry-LaminA-C-18 (Addgene #55068) | pRK5-myc-Miro2 T18N (Addgene #47897) | Addition of mcherry | URDU4 mchr-Miro2 T18N | ATCTCCGAGGAGGACCTGGGATCTATGGTGAGCAAGGGCGAGGAG<br>GATGCGCACGTCCCGCCTCATGCTTCCGCTTCCGCCGG |
| Megaprimer | mCherry-LaminA-C-18 (Addgene #55068) | pRK5-myc-Miro2 ΔTM(Addgene #47901) | Addition of mcherry | URDV3 mchr-Miro2 ΔTM | ATCTCCGAGGAGGACCTGGGATCTATGGTGAGCAAGGGCGAGGAG<br>GATGCGCACGTCCCGCCTCATGCTTCCGCTTCCGCCGG |
| Quickchange | n/a | URDT12 mchr-Miro2 | Mutation of amino acid A13 to V for GTP-bound<br>state | URDW1 mchr-Miro2 A13V | CTTCCCCACCTGGACCTCGCCCAGTAA<br>TTACTGGGCGAGGTCCAGGTGGGGAAG |
| Megaprimer | EGFP-Syne2 (Luxton lab) | URDT12 mchr-Miro2 | Swap of mitochondrial insertion sequence with KASH<br>domain to localize to nuclear membrane | UREL7 mchr-Miro2-KASH | CCCTCTTCCTTCTGGCTCCGGATGCCCCACCTCGACAGCC<br>CAAGTTGGGCCATGGCGGCCTATCTAGACTAGGTGGGAGGTGG |
| Megaprimer | pcDNA3.1-MYO10-HMM-Nanotrap (Addgene #87255) | UREL7 mchr-Miro2-KASH | Swap of Miro2 for an anti-GFP nanobody | URFB5 mchr-Nanotrap$^{green}$-KASH | CATCCGGCGGAAGCGGAAGCAGATATCTGATGGCCCAGGTTCAACT<br>GGCATCCGGAGCCAGAAGGAAGAACTTCCACCTTTAGAGCTCACCGTCACCTGAGT |
| Quickchange | n/a | URDT12 mchr-Miro2 | Mutations to acidic residues in switch I of Miro2<br>N-terminal GTPase | URFF9 mchr-Miro2 switch I mut. | GGGAGGGACCGCCGCGGGGAACGCCGCGCCCACCAG<br>CTGGTGGGCGCGGCGTTCCCCGCGGCGGTCCCTCCC |
| Quickchange | n/a | URFF9 mchr-Miro2 switch I mut. | Mutations to acidic residues in switch II of Miro2<br>N-terminal GTPase | URFJ75 mchr-Miro2 switch I & switch II mut. | GACTACTCAGCAGCCGCGCAGACGGCCGCGGCGCTGCGGGAG<br>CTCCCGCAGCGCCGCGGCCGTCTGCGCGGCTGCTGAGTAGTC |
| Megaprimer | pGEX6P1-mCherry-Nanobody (Addgene #70696) | pEGFP-C1 (Clontech) | Insertion of an anti-mcherry nanobody to the<br>C-terminus of GFP | URET3 GFP-Nanotrap$^{red}$ | CTGCAGTCGACGGTACCGCGATGGCACAGGTTCAGCTGGTTG<br>GTTATCTAGATCCGGTGGATCCCGGTTATGTAAACGGGCTGCTAACGGTAAC |
| Megaprimer | pcDNA3.1-MYO10-HMM-Nanotrap (Addgene #87255) | pGEX6P1-mCherry-Nanobody (Addgene #70696) | Swap of anti-GFP nanobody for an anti-mcherry<br>nanobody (LaM4) | UREK2 MYO10-HMM-Nanotrap$^{red}$ | GAATTCTGCAGATATCTGATGGCCCAGATGGCACAGGTTCAGCTGGTTG<br>CTCTAGACTCGAGCGGCCGCTCATTATGTAAACGGGCTGCTAACGGTAAC |
